# Therapeutic potential of red blood cell-derived extracellular vesicles in reducing neuroinflammation and protecting against retinal degeneration

**DOI:** 10.1101/2024.08.06.606930

**Authors:** Rakshanya Sekar, Adrian V. Cioanca, Yilei (Evelyn) Yang, Karthik Shantharam Kamath, Luke Carroll, Riccardo Natoli, Yvette Wooff

## Abstract

Neuroinflammation is a pathological process mediated through immune cell activation and pro-inflammatory cytokine release, resulting in neuronal cell death. In the central nervous system (CNS), neuroinflammation is a characteristic feature underlying the onset and progression of retinal and neurodegenerative diseases. Targeting neuroinflammation to reduce neuronal cell death and protect against visual and cognitive declines is therefore a key therapeutic strategy. However, due to the complex and multi-faceted nature of these diseases, to date there has been little therapeutic success with single target approaches insufficient to tackle widespread and multi-pathway inflammatory cascades. Furthermore, as the retina and brain reside within immune-privileged environments, a major challenge in treating these diseases is producing and delivering a therapeutic that, in itself, does not exacerbate inflammation. Extracellular vesicles (EV), derived from red blood cells (RBC EV), present a promising solution to overcome these hurdles, due to their innate ability to cross blood-tissue barriers, biocompatible nature, and their broad anti-inflammatory properties to modulate complex neuroinflammatory pathways.

This study therefore investigated the therapeutic potential of RBC EV in mediating neuroinflammation using an *in-vivo* photo-oxidative damage model of retinal degeneration as a model for CNS neuroinflammation. In this work, we developed a novel incubation pipeline using N1 medium supplement and superoxide dismutase (SOD) supplementation to promote the production of safe, neuroprotective, and anti-inflammatory RBC EV. Delivery of RBC EV *in vivo*, was shown to be safe with strong penetration across all retinal layers. Further, therapeutic administration of RBC EV via local intravitreal injection significantly reduced inflammation and cell death and preserved retinal function. Notably, strong safety and therapeutic efficacy was also demonstrated in the retina following systemic (intraperitoneal) administration, highlighting a potential game-changing approach for less-invasive therapeutic delivery to the CNS. Finally, multi-omic analyses and *in vitro* findings supported an anti-inflammatory mechanism-of-action, with RBC EV modulating pro-inflammatory cytokine release, including those known to be involved in the pathogenesis of retinal and neurodegenerative diseases.

Taken together, these findings highlight the broad applicability of RBC EV in treating neuroinflammation in the CNS, presenting a scalable and effective treatment approach for these currently untreatable diseases.

## Introduction

Neuroinflammation within the central nervous system (CNS) plays a key role in the onset and progression of retinal and neurodegenerative diseases [1–3]. While the initial inflammatory response, including the activation of resident immune cells like microglia, aims to protect and repair neural tissue, prolonged immune cell activation and chronic inflammation ultimately results in neuronal cell death and the pathological decline of vision and cognitive functioning [4–6]. Therefore, developing therapeutics that can modulate neuroinflammatory pathways to prevent sustained inflammation and subsequent neuronal cell death holds strong potential for treating both retinal and neurodegenerative conditions [2, 7].

Challenges in developing therapeutics for both retinal and neurodegenerative diseases include overcoming blood-tissue barriers of these immune-privileged environments and modulating complex and multifaceted neuroinflammatory pathways [7–9]. The blood-brain barrier (BBB) and blood-retina barrier (BRB) exclude most therapeutics following systemic administration, necessitating invasive delivery methods [10]. These drugs are therefore often also encapsulated within synthetic nanoparticles, which can induce neurotoxicity [11] and provoke additional pro-inflammatory responses [12, 13]. Moreover, many drugs in development and clinical trials are designed to target single inflammatory pathways [14–16], which may be insufficient to address the multiple pathways involved in complex neurodegenerations such as Alzheimer’s disease, Parkinson’s disease and age-related macular degeneration (AMD) - a leading cause of blindness in the world [17]. As AMD alone affects 1:7 people over the age of 50, or 200M worldwide [18] with no widely available treatments [15, 16], there is an urgent need to develop broadly anti-inflammatory and neuroprotective therapeutics to treat this, and other retinal and neurodegenerative diseases.

Extracellular vesicles (EV) offer a promising solution to overcome many of these challenges. Previously thought to be cellular debris, EV are now recognized as key players in both normal physiological processes and pathological conditions [19–22]. EV are ubiquitously distributed in bodily fluids and play essential roles in intercellular communication by transporting bioactive molecules to adjacent and distant cell types to regulate biological pathways [23–25]. In fact, EV-mediated transport of molecular cargo has been shown in both the retina, and CNS to modulate inflammation across health and disease [20, 26–31]. Unlike synthetic nanoparticles, EV are cell-derived and offer higher biocompatibility [32], better biodistribution profiles [33], and low immunogenicity [34]; making them ideal therapeutic agents for treating neuroinflammation.

Red blood cell (RBC) derived EV are particularly ideal due to their inherent safety profile, homogeneity, and innate anti-inflammatory properties [35–37]. As mature RBCs do not have nuclear or mitochondrial DNA, RBC EV are less likely to carry bioactive molecules, reducing horizontal transfer [38]. With their abundant and homogenous nature, RBC are easy to collect and process in large quantities [39, 40], overcoming significant manufacturing hurdles associated with scale, homogeneity, and batch reproducibility as commonly seen with stem cell-based EV production [41]. Importantly, RBC EV express CD47 on their surface, inhibiting phagocytosis through interaction with the macrophage inhibitory receptor signal regulatory protein alpha (SIRPα), thus reducing RBC EV clearance and reducing adverse inflammatory reactions [42–45]. Notably, RBC EV have been shown to intrinsically induce anti-inflammatory effects in macrophages further enhancing their suitability for clinical applications in treating diseases of the brain and retina [40, 46]. The natural anti-inflammatory properties and clinical viability of RBC EV makes them ideal neuroinflammatory modulators for the CNS, which could be leveraged for the development of effective and biocompatible therapies for treating neurodegenerative disorders where neuroinflammation is an underlying feature, including AMD.

This study therefore aims to investigate the therapeutic potential of RBC EV in mediating neuroinflammation in the retina using an *in-vivo* photo-oxidative damage induced model of retinal degeneration [47]. The retina, as an extension of the CNS, displays similar anatomical, functional, and immunological aspects to the brain offering a valuable opportunity to study shared clinical and pathological features [48]. We have previously shown that mice undergoing photo-oxidative damage have increased levels of inflammation, initiating pathological changes such as immune cell activation and recruitment, and focal photoreceptor cell death [47] - hallmark features of currently incurable retinal degenerative diseases such as AMD.

In this study, we pioneered a unique incubation strategy to produce high yields of reproducible anti-inflammatory and neuroprotective RBC EV, using N1 medium supplement and superoxide dismutase (SOD). *In vivo* delivery of these RBC EV (N1+SOD), both locally and systemically, was not only found to be safe, but showed strong therapeutic efficacy in protecting against retinal degeneration, with RBC EV (N1+SOD) treated mice exhibiting reduced inflammation, decreased retinal cell death, and preserved retinal function. Furthermore, using *in vitro* assays and multi-omics analyses, we demonstrated that RBC EV (N1+SOD) were able to modulate the release of key pro-inflammatory cytokines known to be involved in retinal degenerations, such as IL-6 and IL-1β, supporting an anti-inflammatory mechanism-of-action likely contributing to the observed retinal protection. These results combined supports the development of RBC EV for the treatment of retinal degenerations and suggests broader therapeutic applicability of RBC EV in modulating inflammation in the CNS, presenting a scalable, effective, and potentially less-invasive systemic treatment approach for wider neurodegenerative conditions.

## Methods

### 2.1 Red blood cell (RBC) processing and characterisation

#### 2.1.1 RBC processing

Whole mouse blood in EDTA was purchased from Applied Biological Products Management (MSBX 0005 – 5ml tubes). Whole blood was spun at 600 xg for 20 minutes to separate the plasma, buffy coat, and RBC. The plasma and buffy coat were removed via pipetting, and the remaining RBC were diluted 1:10 in 1X PBS (Gibco; pH 7.2). RBC suspension was passed through a leukocyte depletion filter (Sterile Acrodisc® WBC syringe filter with Leukosorb Membrane, 25 mm; Pall) to remove contaminant leukocytes.

RBC suspension was incubated in T25 flasks supplemented with either: N1 Medium Supplement (100X, Sigma; N6530) at 1:100 dilution and/or superoxide dismutase (SOD) (Worthington Biochemicals; LS003540) at 5 ug/ml, Trans-resveratrol (Sigma PHR2201-200MG), Kaempferol (Sigma, 60010-25MG) or L-Glutathione reduced (Sigma, G4251-1G) for 18 hours at 37 °C on a shaker incubator at 90 rpm.

#### 2.1.2 RBC health measurements

Post incubation, the RBC suspension was transferred to 15 mL falcon tubes and spun at 600xg for 20 minutes to separate RBC from extracellular vesicles (EV) in suspension [36]. To determine the effect of incubation on the health of RBC, the 600 xg pellet was resuspended in 1X PBS (Gibco; pH 7.2) in 1:1000-1:10000 dilution range to perform qualitative and quantitative assessments. 200 µL of pellet suspension at 1:1000 was applied to a glass hemocytometer (Westlab, product number 071301-9877) and the number of RBC in the large centre square were counted to determine total cell number. Zeiss Axiovert 200 microscope was used to take images of RBC at 4x and 20x magnification and images were processed using ImageJ software (version 2.1.0). RBC abnormality percentage was determined by studying RBC membrane and classifying them based on size (normal, microcyte and macrocyte) and membrane shape variation (normal, burr cell, tear drop, sickle cell, ovalocyte and schistocyte), all statistical analysis was performed on Prism 9.

#### 2.1.3 RBC viability measurements

To determine the effect of supplementation on RBC viability post incubation, Calcein-AM (C1430, Thermo Fisher Scientific) and Annexin V (563973, BD Biosciences) staining was performed and assessed using imaging flow cytometry (ImageStream®X MkII). Calcein-AM was prepared as a stock solution of 10 mM in dimethyl sulfoxide (DMSO) and a working solution of 100 μM in PBS buffer, pH 7.4 was prepared as required. 2 × 10^5^ RBCs in 200 μl PBS, were incubated for 45 minutes with Calcein-AM working solution (final concentration in Calcein-AM, 5 μM) at 37°C in the dark to preserve fluorescence [49]. Cells were then spun down by centrifugation (1000 xg at 4°C for 5 min), resuspended in 0.5 ml of Annexin V binding buffer containing 2 mM calcium chloride and incubated with 5 μl Bv421-Annexin-V for 15 min at room temperature in the dark. Cells were analysed for biparametric histograms FL1 (Calcein-AM) versus FL2 (Annexin-V) using ImageStreamX Mk II (Amnis Corporation, Seattle, WA, USA). RBC experimental permeabilization as a control of dying cells was readily made with 1% saponin [50].

### 2.2 RBC EV processing

#### 2.2.1 RBC EV isolation

Following the 600 xg spin to remove RBC, the RBC supernatant was transferred to 15mL falcon tubes and spun at 1500 xg for 15 minutes and 3200 xg for 10 minutes, excluding the pellet at each spin step by transferring the supernatant to a fresh tube. Following, RBC supernatant S1 was transferred to ultracentrifuge tubes (Ultra-Clear Thinwall Tubes 13.2ml, Beckman Coulter, USA) and spun at 10,000 xg for 35 minutes. The resultant pellet P2 was discarded and the supernatant S2 was transferred to fresh ultracentrifuge tube and re-spun at 100,000 xg for 1h 30 minutes. P3 containing RBC EV was then collected, washed in PBS via tituration, and re-spun at 100,000 xg for 1h 35minutes. Finally, P4 (RBC EV pellet) was resuspended in 50-500uL (depending on downstream experiment) and/or snap-frozen at −80C until use.

#### 2.2.2 RBC EV characterisation

##### NanoSight characterisation

The concentration and size distribution of RBC EV was measured using nanoparticle tracking analysis on a NanoSight NS300 (Malvern Instruments, Malvern, United Kingdom) and/or ZetaView® QUATT (Particle Metrix, Ammersee, Germany) instrument. RBC EV were diluted 1:10,000 in 1x PBS to achieve a particle per frame value between 20 and 100. The mean concentration value for each sample, in particles/mL, was exported to Prism 9 for plotting and statistical analyses [20].

##### Cryogenic electron microscopy

RBC EV samples (1:1000) in 1x PBS were applied to a glow-discharged (PELCO easiGlow™ Glow Discharge Cleaning System) 300 mesh EM grid with lacey carbon for vitrification at 80-90% humidity, in room temperature. Any excess sample present was removed by blotting with filter paper. Loaded grids were placed into liquid ethane (kept in equilibrium with solid ethane) using a Leica EMGP2 Automatic Plunge Freezer (Centre for Advanced Microscope facility, JCSMR) [51]. This grid was then stored in liquid nitrogen until further use. A JEOL JEM-F200 microscope was used to image EV at 200kV voltage and 16,000x magnification. Images were processed using ImageJ. All EV visible across 10 images were measured for size (in nanometres) and exported to Prism 9 for plotting and statistical analysis.

##### Western blot analysis

Total lysates were extracted from cells and RBC EV by incubating with RIPA buffer supplemented with 1:100 protease inhibitor cocktail (Sigma-Aldrich, MO, United States). 10-20 ug of protein lysates/well were separated on 4-12% polyacrylamide gels (Thermo Fisher Scientific, MA, United States) at 100 V for 60 minutes and transferred to a Nitrocellulose membrane (Bio-Rad, CA, United States) using a Power Blotter semi-dry system (Thermo Fisher Scientific, MA, United States) at 20V for 15 minutes. Membranes were washed in PBS-Tween (0.01%; PBS-T), blocked in Pierce™ Clear Milk Blocking Buffer (Thermo Fisher Scientific, MA, United States) for 1 hour and then incubated overnight at 4°C with primary antibodies TSG101 (1:1000, ab30871, Abcam, Cambridge, United Kingdom), ALIX (1:1000, EPR23653-32, ab275377,Abcam, Cambridge, United Kingdom), Calnexin; CALX (1:1000, ab22595, Abcam, Cambridge, United Kingdom) or GAPDH (1:2000, G9545-100UL, Sigma-Aldrich, United States). Following three washes in PBS-T, blots were incubated in appropriate secondary antibodies, HRP-conjugated Goat Anti-Rabbit IgG (H + L) (1:1000, 170-6515, Bio-Rad, CA, United States) or Goat-anti-Mouse IgG (1:1000, 170-6516, Bio-Rad, CA, United States) for 2 h at room temperature. Membranes were washed in PBS-T and developed for 2 min with ClarityTM Western ECL Substrate (Bio-Rad, CA, United States). Imaging was performed using a ChemiDocTM MP Imaging System with Image LabTM software (Bio-Rad, CA, United States).

#### 2.2.3 RBC EV labelling

To determine RBC EV uptake *in vivo* and *in vitro*, RBC EV were labelled using the SYTO^TM^ RNASelect^TM^ Green Fluorescent Cell Stain, a cell-permeant RNA-specific stain. The stain was first diluted 1:5 in DMSO (#ICN19141880, Thermo Fisher Scientific, MA, United States) to give a 1mM DMSO stock. The RBC EV stock was diluted in sterile 1x PBS to a total volume of 100µL. 1x PBS was used as a control for background staining. 1µL of the DMSO stock was then added to the diluted RBC EV, mixed thoroughly, and incubated at 37 °C for 30 minutes (dark). At approximately halfway, the solutions were mixed to ensure the dye did not settle at the bottom of the tube. Afterwards, the solutions were added to Amicon^TM^ 50kDa MWCO filter units (UFC505024, Merck) pre-rinsed with UltraPure^TM^ distilled water, and spun at 5000xg for 20 minutes to remove excess unbound dye. The solutions were further washed thrice by adding 500µL of sterile 1x PBS and spinning the columns at 10,000xg for 10 minutes each time. The solutions were then made up to the same volume using sterile 1x PBS. The fluorescently labelled RBC EV or PBS-only control solutions were then used to determine RBC EV uptake *in vivo*.

#### 2.2.4 RBC EV cell viability and inflammatory profiling

RBC EV (N1+SOD) in frozen −80 C aliquots were sent to CruxBioLabs (VIC, AUS) for independent validation of safety and anti-inflammatory properties using control and 100 ng/mL LPS (L6529-1MG, Thermo Fisher Scientific, MA,USA) stimulated human peripheral bone mononuclear phagocyte cells (50,000 PBMCs). Procartaplex (Thermo Fisher Scientific, MA, USA) custom panel 8-plex for IL-1α, IL-1β, IL-6, IL-8, IL-10, MCP-1, MIP-1α, TNFα according to the manufacturer’s instructions (Cat# MCYTOMAG-70K, Merck Millipore, MA, USA).

### 2.3 Quantitative Proteomics on RBC and RBC EV

To characterize the proteomic signature of RBC and RBC EV with or without N1 and/or SOD supplementation, snap-frozen cell and EV pellets were used and processed by Australian Proteome Analysis Facility (APAF), Macquarie University, NSW, Australia. Protein concentrations were determined using a BCA assay. Fifty micrograms of protein from each sample were aliquoted and volumes were normalized with 100 mM TEAB and 1% SDC. Disulfide bonds were reduced with 10 mM DTT at 60 °C for 30 minutes and alkylated with 20 mM IAA at room temperature in the dark for 30 minutes. Proteins were digested with trypsin at a 1:25 enzyme-to-protein ratio for 16 hours at 37 °C. Subsequently, samples were acidified to 1% formic acid, centrifuged to pellet SDC, and the supernatant was desalted using Stage-Tip purification and dried via vacuum centrifugation. Reconstituted samples were diluted with 0.1% formic acid for LC-MS/MS analysis.

For ion-library creation using high pH RP-HPLC fractionation, a small portion of peptides from each sample were pooled, desalted using a C18 column, and dried. The peptide mixture was resuspended in 5 mM ammonia solution (pH 10.5), fractionated using an Agilent 1260 HPLC system over a 55-minute gradient, and consolidated into 20 fractions. Dried fractions were reconstituted in 0.1% formic acid for LC-MS/MS analysis.

LC-MS/MS analysis was performed using a Thermo Orbitrap Exploris mass spectrometer coupled with a Vanquish Neo UHPLC system. Peptides were trapped on a C18 PepMap 100 column and separated on a 75 µm × 30 cm C18 analytical column at 45 °C. The mobile phase consisted of 99.9% water with 0.1% formic acid (A) and 80% acetonitrile with 0.1% formic acid (B). Peptides were eluted over a linear gradient from 2.5% to 37.5% mobile phase B for 59 minutes at a flow rate of 300 nL/min. For Data-Dependent Acquisition (DDA) of HpH-fractionated samples, peptide precursors (350-1850 m/z) were scanned at 60k resolution with a 1.5 s cycle time and dynamic exclusion set to 15 seconds. Fragmentation was carried out using HCD at 27% normalized collision energy, and fragment ions were detected at 15k resolution. For Data-Independent Acquisition (DIA), peptides were eluted similarly, with MS1 scans at 60k resolution (350-1450 m/z) and MS2 scans at 30k resolution using 20 variable windows and 27% collision energy.

Data analysis was performed as previously described [27]. Raw DDA data were processed with FragPipe version 20.0 and MSFragger version 3.8 using a FASTA file containing *Mus musculus*, Bovine SOD, and decoy protein sequences (https://www.uniprot.org/, 55,022 sequences, 25-Aug-2023). Parameters included trypsin (semi-specific) with two missed cleavages, 20 ppm precursor and fragment mass tolerances, oxidation (M) as a dynamic modification, and carbamidomethyl (C) as a static modification. DIA data files were processed using DIA-NN version 1.8 with default parameters, using ion-library data generated from FragPipe. All DIA data files were loaded into DIA-NN, with output filtered at 0.01 FDR. Library precursors were reannotated using the FASTA database, enabling N-terminal methionine excision. Tryptic peptide digestion allowed a maximum of two missed cleavages. Peptide length was set between 7 and 30 residues, and precursor m/z ranged from 300 to 1800, with charge states between 1 and 4. Cysteine carbamidomethylation was a fixed modification, with a maximum of one variable modification.

#### 2.3.2 Data analysis

Data analysis was performed as described previously [27]. The initial processing of protein intensity data began with the application of a base-2 logarithmic transformation excluding proteins with more than two missing values, within any single experimental group to maintain data integrity. For proteins with missing values, imputation strategy was employed. Post-imputation, data normalization was performed using variance stabilization transformation (VST) via the normalize_vst function in the R DEP package, ensuring homoskedasticity. The processed data were then subjected to differential expression analysis following the limma pipeline, a well-established methodology in the field of bioinformatics for identifying differentially expressed genes or proteins with precision and reliability. Pathway analysis was conducted through gene set enrichment analysis (GSEA) utilizing the fgsea function using C2 canonical and C5 gene ontology database from the Molecular Signatures Database (MsigDB) serving as the reference gene sets.

Data tables for RBC and RBC-EV proteomics can be found in *Supplementary Table 1*

### 2.4 Gene Expression Analysis

#### 2.4.1 RNA extraction

Small RNA extraction began with the lysis of RBC EV in 300 μL of lysis buffer, followed by the addition of 30 μL miRNA homogenate, and incubated on ice for 10 minutes. Subsequently, 330 μL Acid-Phenol:Chloroform was added, and the mixture was vortexed for 60 seconds. Centrifugation at 10,000xg for 5 minutes facilitated phase separation. The aqueous phase was transferred and mixed with 1.25x volume of ice-cold absolute ethanol for total RNA isolation. The mixture was then passed through a purification column. The column was washed with 700 μL miRNA wash solution #1 and twice with 500 μL wash solution #2, then spun dry. RNA was eluted with heated solution (95°C), collected into 30 μL, and stored at −80°C. RNA quality and concentration were measured using a Nanodrop ND-1000 spectrophotometer and Agilent 2100 Bioanalyzer.

#### 2.4.2 Library preparation and sequencing

The library preparation for high-throughput sequencing of RNA from RBC EV was carried out at the Biomolecular Research Facility (JCSMR, ANU). Libraries were synthesized using the Capture and Amplification by Tailing and Switching (Diagenode). Libraries were sequenced on the Illumina NovaSeq 6000 acquiring at least 10 million 50bp single-end reads per sample as described previously [20].

#### 2.4.3 Bioinformatics

Data analysis was performed as described in previous publication [20, 52]. For preprocessing-indexes, adapters, and template-switching oligonucleotides were removed with *cutadapt.* Subread-align was used of reads to the mm10 mouse genome following which *featureCounts* was run for quantifying miRNA alignments. The following command sequence voom --> lmFit --> ebayes --> topTable was used for assessing differential gene expression data in Rstudio using limma package.

The RNA data that support the findings of this study are openly available in the Sequencing Read Archive NCBI repository under bioproject reference number PRJNA1144155.

### 2.5 Animal Handling and Paradigms

All experiments were conducted in accordance with the ARVO Statement for the Use of Animals in Ophthalmic and Vision Research and with approval from the Australian National University’s (ANU) Animal Experimentation Ethics Committee (AEEC) (Ethics ID: A2020/41 and A2023/318; Rodent models and treatments for retinal degenerations). Adult male and female C57BL/6J wild-type (WT) mice (aged 60-80 postnatal days) were purchased from the Animal Resources Centre (ARC, Canning Vale, Western Australia (WA)). Mice were bred, reared, and housed under 12 h light/dark cycle conditions (5 lux) with free access to food and water.

Photo-oxidative damage (PD): Animals in the photo-oxidative damage group (PD) were placed into Perspex boxes coated with reflective interior surface and exposed to 100 K lux white light from light-emitting diodes (LED) for a period of 5 days and were administered a pupil dilator (Minims® atropine sulphate 1% w/v; Bausch and Lomb) to both eyes twice a day (9am and 4 pm) during the damage paradigm.

Dim-reared (DR) : Dim-reared control mice were maintained in standard housing conditions of 12 h light (5 lux)/dark cycle with free access to food and water.

Age-matched and sex-matched mice were randomly assigned to 5-day photo-oxidative damage (PD) or dim-reared control (DR) groups (N = 5-15 mice per group, each eye measured independently as biological replicate).

### 2.6 *In vivo* administration of RBC EV

To test safety and therapeutic efficacy of RBC EV, mice were administered with RBC EV both locally and systemically in dim-reared (DR) or photo-oxidative damage (PD) conditions. Mice were either injected with RBC EV (N1+SOD), RBC EV (SOD), RBC EV (N1), RBC EV (PBS), or PBS (N1+SOD), PBS (SOD), PBS (N1) or PBS as controls. In some experiments, RBC EV or PBS were labelled with SYTO RNASelect to fluorescently label EV RNA as per above method. Mice from the same litter were used in each experiment for comparison and were randomly assigned across treatment groups. Retinal function and morphological assessments were performed using ERG, OCT and histology as listed in *2.5 Retinal Assessment*.

#### 2.6.1 Intravitreal injections

Mice were anaesthetised with an intraperitoneal injection of Ketamine (100mg/kg) and Xylazil (10mg/kg) and then 1% w/v Atropine was applied to the ocular surface of each eye to dilate the pupils. Intravitreal injections were performed as described in detail previously [53]. A 10µL NanoFil syringe, with a 34G needle was used to inject 1µL (2.0×10^9^ EV) of each EV mixture or control solution through the previously made pilot hole into the vitreous of the eye. After injection, the eye was swabbed with 1% Chlorsig to prevent bacterial infection and GenTeal^TM^ Gel to prevent eye dryness. To aid recovery, animals were intraperitoneally injected with Reversamed (1mg/kg).

Animals were placed into photo-oxidative damage for 5 days (degeneration paradigm) or were kept in standard housing conditions for 7 days (safety assessment).

#### 2.6.2 Intraperitoneal injections

Mice were injected via intraperitoneal injection at 2.0×10^11^ EV/100µL dose daily for 5 days (photo-oxidative damage) or were kept in standard housing conditions for 7 days (dim-reared safety assessment).

### 2.7 Retinal Assessment

#### 2.7.1 Retinal function via electroretinography (ERG)

Full-field scotopic electroretinography (ERG) was performed to assess the retinal function of control or treated mice in DR conditions or following PD as described [54]. Mice were dark-adapted overnight before being anaesthetised with an intraperitoneal injection of Ketamine (100 mg/kg; Troy Laboratories, NSW, Australia) and Xylazil (6 mg/kg; Troy Laboratories, NSW, Australia) diluted in PBS. Both pupils were dilated with one drop each of 2.5% w/v phenylephrine hydrochloride and 1% w/v tropicamide (Bausch and Lomb, NY, USA).

Anaesthetised and pupil dilated mice (as above) were placed on the thermally regulated stage of the Celeris ERG system (Diagnosys LLC, MA, USA) and response from both eyes was simultaneously recorded as previously described [54]. Responses were recorded and analysed using Espion V6 Software, (Diagnosys LLC, MA, USA). Statistics were performed in Prism V9.4 (GraphPad Software, CA, USA) using a two-way analysis of variance (ANOVA) to test for differences in a-wave and b-wave responses. Data were expressed as the mean wave amplitude ± SEM.

#### 2.7.2 Retinal tissue collection and preparation

Animals were euthanised with CO2 following functional assessment. The superior surface of the right eye from each animal was marked and enucleated, then immersed in 4% paraformaldehyde (PFA) in PBS for 3 h at 4 °C. Eyes were then cryopreserved in 15% sucrose solution overnight, embedded in OCT medium (Tissue Tek, Sakura, JP) and cryosectioned at 12 μm in a parasagittal plane (superior to inferior) using a CM 1850 Cryostat (Leica Biosystems, Germany) [47, 54]. The retina from the left eye of each mouse was excised through a corneal incision and placed into 300uL of Lysis/Binding Buffer (Thermo Fisher Scientific, MA, USA) and stored at − 80 °C until further use.

#### 2.7.3 TUNEL assay

Terminal deoxynucleotidyl transferase (Tdt) dUTP nick end labelling (TUNEL) was used as a measure of photoreceptor cell death. TUNEL in situ labelling was performed on cryosections using a Tdt enzyme (Cat #3,333,566,001, Sigma-Aldrich, MO, USA) and biotinylated deoxyuridine triphosphate (dUTP) (Cat #11,093,070,910, Sigma-Aldrich, MO, USA) as previously described [47, 54]. Images of TUNEL stained sections were captured with the A1 + confocal microscope at 20× magnification.

#### 2.7.4 IBA-1 immunohistochemistry

Immunolabeling for IBA-1 (1:1000, Anti-Iba1, Rabbit, 019-19741, Wako, Osaka, Japan) and quantification was performed as previously described [47]. The number of IBA-1+ cells (a marker of retinal microglia and macrophages) was counted across the superior and inferior retina using two retinal sections per mouse and then averaged. Retinal cryosections were stained with the DNA-specific dye bisbenzimide (1:10,000, Sigma-Aldrich, MO, United States) to visualize the cellular layers. Images of IBA-1 + staining were captured with the A1 + confocal microscope at 20× magnification.

### 2.8 Imaging and statistical analysis

Morphology of RBC was examined using Zeiss Axiovert 200 Fluorescence/Live cell imagine microscope (Zeiss). Fluorescence in retinal sections was visualised using A1+ Confocal Microscope System with NIS-Elements Advanced Research software.

RNA and Proteomic analysis: Statistical analysis for multiomics was performed as specified above. Graphing was generated using ggplot2 package and Prism V9.0.

Functional analysis (ERG): All graphing and statistical analysis was performed using Prism 9 (GraphPad Software, CA, USA). Two-way analysis of variance (ANOVA) with Šídák or Bonferroni multiple comparison post-hoc test was used to determine the statistical outcome; p value of < 0.05 was considered statistically significant. All data were expressed as the mean ± SEM.

Histology: All graphing and statistical analysis was performed using Prism 9 (GraphPad Software, CA, USA). An unpaired Student t test, one-way analysis of variance (ANOVA), or two-way ANOVA with Tukey’s multiple comparison post-test were utilised to determine the statistical outcome; p value of < 0.05 was considered statistically significant for all histological analysis. All data were expressed as the mean ± SEM. For TUNEL and IBA counts, 2 sections per mouse were used. If any section was partially missing due to cryosectioning, it was excluded from the analysis.

*In vitro* experiments: All graphing and statistical analysis was performed using Prism 9 (GraphPad Software, CA, USA). Unpaired t test, Mann–Whitney test with false discovery rate (FDR), or two-way ANOVA with Tukey’s or Šídák multiple comparison post-hoc test was used to determine the statistical outcome. p value of < 0.05 was considered statistically significant.

*In vivo* experiments: ‘N’ represents the number of animals in the individual experimental run, graphs depict average of technical replicates from multiple measurements per animal.

## Results

### Result 1: Media supplementation influences red blood cell (RBC) viability, profile, and proteomic composition

To identify the optimal media supplementation for maintaining red blood cell (RBC) integrity, RBC were isolated following plasma and leukocyte depletion and incubated overnight at 37°C in 1:10 PBS, with or without the addition of culture supplements: N1 media supplement, Superoxide Dismutase (SOD), N1+SOD, Resveratrol, Kaempferol, and Glutathione (**Figure 1A**). Light microscopy was used to evaluate the membrane abnormalities of RBC post-treatment to determine optimal supplementation dosage. Dilution curve analyses identified the optimal supplementation dosage resulting in minimal membrane abnormalities: Resveratrol at 100 µM, Kaempferol at 2 µM, and Glutathione at 0.1 mM **(Supplementary Figure 1A, P<0.05)**. These dosages were then re-evaluated against pre-treatment (Pre-Tx), PBS-only control, N1 media supplement (1X), and/or SOD (5ug/mL). Resveratrol (100 µM), N1 media supplement (1X) and/or SOD (5ug/mL) significantly reduced membrane abnormalities compared to both pre-treatment and PBS-only control (**Figure 1B, P<0.05)**. Furthermore, RBC counts were performed to quantify the preservation of cells post-incubation, with the RBC (N1+SOD) group indicating the highest preservation and the RBC (Resveratrol, 100uM) group the least (**Figure 1C, P<0.05),** compared to Pre-Tx. Hence, subsequent experiments progressed with supplements N1 and/or SOD only.

**Figure 1:**
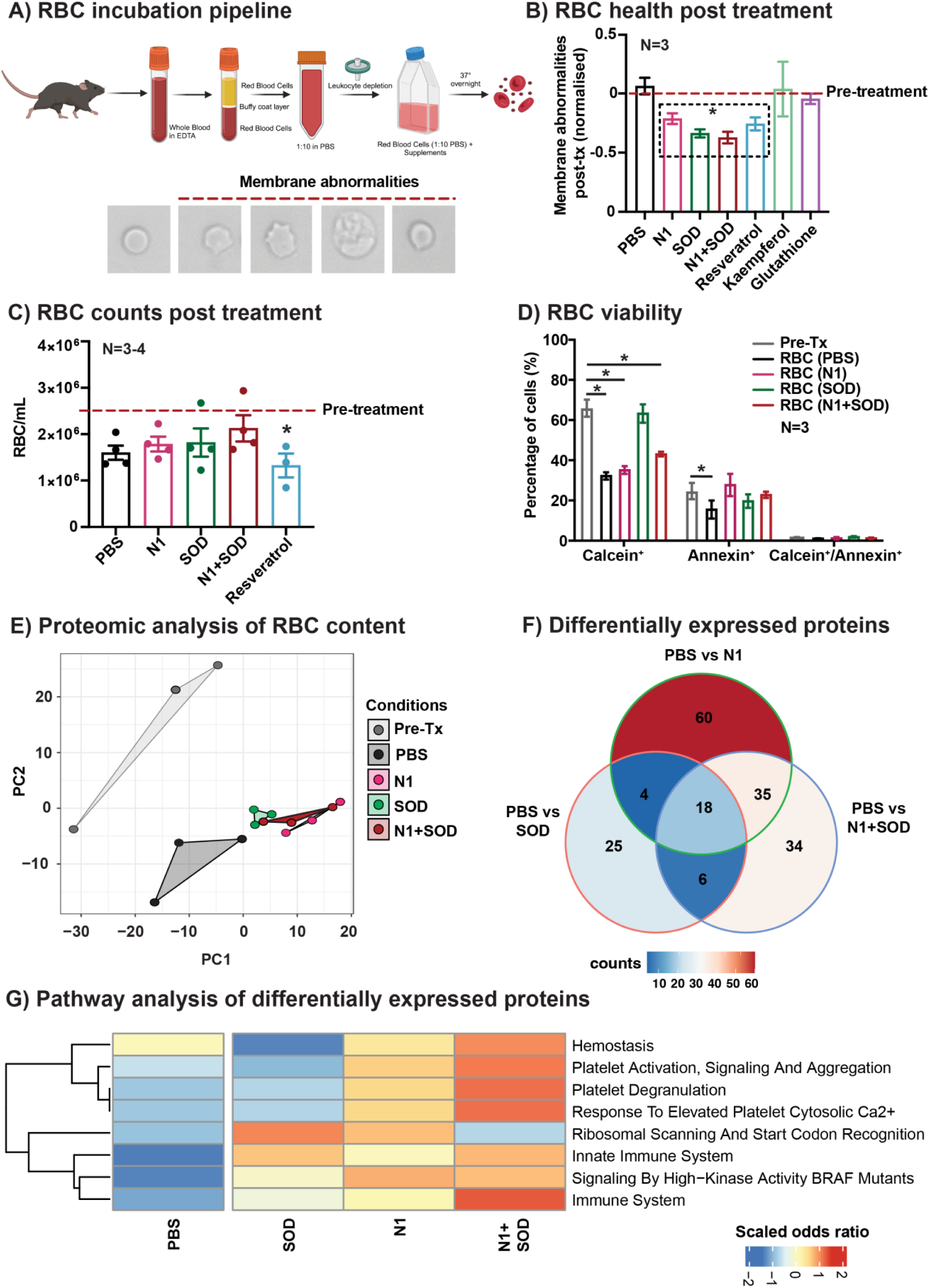
Effects of media supplementation on RBC health, viability, and proteomic content. **(A)** Experimental paradigm: RBC population enriched from whole blood (EDTA) following centrifugation and leukocyte depletion, then incubated in 1:10 PBS with supplements**. (B)** RBC membrane abnormalities post-treatment compared to Pre-Tx control as measured by light microscopy (N=3, P<0.05) **(C)** RBC counts post-treatment compared to Pre-Tx control as measured by hemocytometer (N=3-4, P<0.05**) (D)** RBC viability post-treatment compared to Pre-Tx control as measured using Amnis II imaging flow cytometry (N=3, P<0.05). The proteomic content of RBC analysed using 1D LC-MS/MS, following which **(E)** PCA showed a large variation between treated groups, in particular, compared to Pre-Tx samples. **(F)** Venn diagram shows uniquely expressed proteins in supplement groups compared to relative difference between PBS and Pre-Tx. **(G)** Heatmap displaying pathways associated with proteins enriched in RBC groups compared to PBS controls (N=3).

To elucidate the effects of N1 and/or SOD on RBC viability, imaging flow cytometry was performed to quantify the percentage of Calcein-positive (viability marker), Annexin-positive (apoptosis marker), and double-positive cells. RBC treated with SOD alone displayed comparable viability to that of Pre-Tx, followed by RBC treated with N1+SOD (**Figure 1D, P<0.05)**. The gating strategy and representative images for imaging flow cytometry are presented in **Supplementary Figure 1B-C**.

Following, the proteomic content of RBC (post-treatment) was analysed comparing between supplement groups, PBS-only, and Pre-Tx control. PCA biplots, derived using the protein expression values of all identified proteins, showed a large variation between treated groups, in particular, compared to Pre-Tx samples (**Figure 1E, Supplementary Figure 2A-2B).** It was also demonstrated that RBC with supplements each had uniquely expressed proteomic composition, with RBC (N1) showing the highest number of uniquely expressed proteins compared to RBC (PBS) samples (**Figure 1F**). Finally, for pathway analysis of differentially expressed proteins, PBS was first compared to Pre-Tx control, and this relative difference was compared to each supplement group, identifying that proteins enriched in RBC (N1+SOD) were involved in homeostasis, platelet activation, coagulation, and immune system regulation, including innate immune system regulation (**Figure 1G).**

### Result 2: Media Supplementation modulates proteomic signature and anti-inflammatory properties of red blood cell-derived extracellular vesicles (RBC EV)

Having demonstrated that supplementation can alter RBC health, viability, and proteomic content, we investigated whether the N1 and/or SOD supplementation(s) have subsequent effects on the RBC EV profile and molecular composition. Following overnight incubation, the cell-free supernatant was processed using differential ultracentrifugation to obtain RBC EV (**Figure 2A and Supplementary Figure 3A).** EV were then characterised using nanoparticle tracking analysis which showed comparable size distribution profiles ranging between 100-400 nm (**Figure 2B**) irrespective of supplementation conditions. There was no significant change in mean (**Figure 2Ci**) and mode (**Figure 2Cii**) size of RBC EV between the supplementation conditions, however RBC EV (N1+SOD) generated significantly higher number of EV in comparison to RBC EV (PBS), RBC EV (N1) and RBC EV (SOD) treatments (**Figure 2Ciii**). RBC EV (N1+SOD) were further characterised using cryogenic electron microscopy for size and morphology **(Supplementary Figure 3B and 3C).** Western blotting was performed to detect markers Calnexin (CNX), Tumor susceptibility gene 101 (TSG101) and ALIX (100 kDa). The CNX band (90 kDa) was observed in retinal lysates but was absent in RBC EV lysates, indicating no cell contamination. **(Supplementary Figure 3D).** EV markers ALIX were enriched in RBC EV fractions in comparison to RBC lysates **(Supplementary Figure 3E).** Multiple batches of RBC EV were tested for reproducibility by quantifying size and profile using NTA **(Supplementary Figure 3F).**

**Figure 2:**
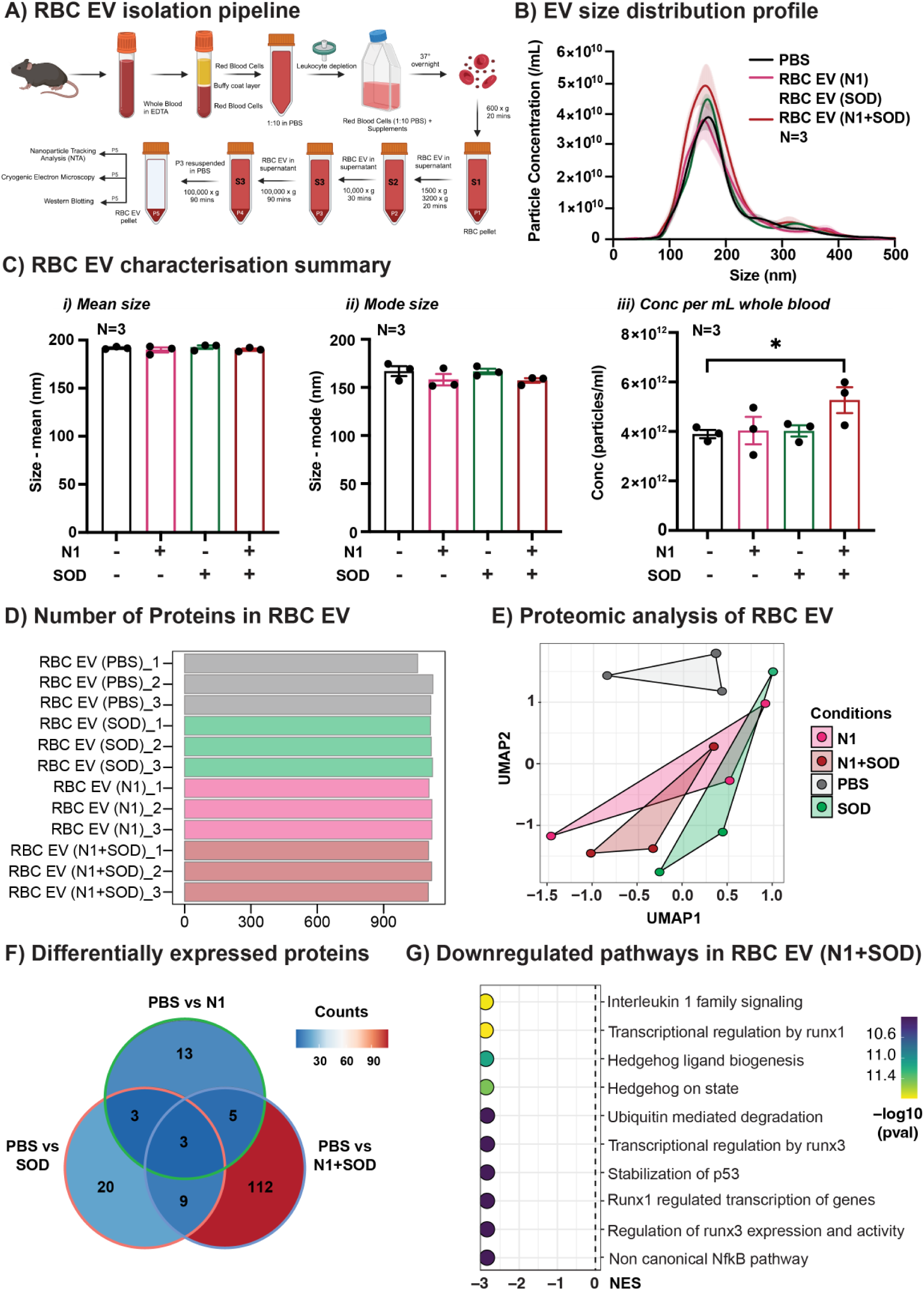
Effects of media supplementation on RBC EV profile and proteomic composition. **(A)** Experimental paradigm. RBC EV populations were isolated post-treatment using differential ultracentrifugation. **(B)** RBC EV size distribution profile post-treatment as measured using NTA (N=3)**. (C)** RBC EV (i) mean size, (ii) mode size and (iii) particle concentration/mL of whole blood as measured using NTA (N=3, P<0.05). Proteomic profile of RBC EV was examined using label-free 2D LC-MS/MS which showed **(D)** comparable numbers of proteins between treatment groups, but **(E)** distinct clustering between sample groups **(F)** Venn diagram represented uniquely expressed proteins within each group, highlighting the most numerous unique proteins were found in RBC EV (N1+SOD). **(G)** Pathway analysis identified that significantly downregulated proteins in RBC EV (N1+SOD) were associated with immune processes and transcriptional regulation (N=3, P<0.05).

Following characterisation, the proteomic composition of RBC EV was also explored. The analysis revealed a comparable number of proteins across all groups – RBC EV (PBS), RBC EV (N1), RBC EV (SOD), and RBC EV (N1+SOD) as shown in **Figure 2D**. However, variations in protein composition were observed, with the RBC EV (N1+SOD) group displaying the most variable expression profile compared to RBC EV (PBS), RBC EV (N1), and RBC EV (SOD) groups (**Figure 2E**). This was also reflected in the high number of unique proteins found within this group (**Figure 2F**). Pathway analysis of differentially expressed (DE) proteins between these groups showed that, in the RBC EV (N1+SOD) group, downregulated proteins were associated with several inflammatory pathways. These included ‘interleukin 1 family signalling’ and the ‘non-canonical NfκB pathway,’ as well as pathways involved in the transcriptional regulation of inflammatory, apoptotic, mitotic, and hematopoietic processes (**Figure 2G).**

Finally, small RNA sequencing was performed on RBC EV (PBS, N1, SOD and N1+SOD) groups to identify if there were any molecular changes in the microRNA signature as a response to supplementation. Out of the 260 total microRNA detected, it was identified that in all samples the top 11 microRNA were highly abundant and made up a significant fraction of the total reads, with RBC-enriched microRNA miR-451 found to be the most abundant **(Supplementary Figure 4A).** Pathway analysis of known targets of the top 11 microRNA identified regulation of processes such as transcription, biogenesis, and senescence **(Supplementary Figure 4B).** Venn diagram showed that there were no unique proteins between groups, suggesting that supplementation did not cause any changes to the microRNA signature of RBC EV **(Supplementary Figure 4C).**

Overall, these results support that media supplementation significantly affects the RBC EV profile and proteomic signature. Specifically, N1+SOD supplementation results in a higher EV concentration and distinct proteomic changes indicating an anti-inflammatory phenotype, more pronounced than other supplementation groups and controls, highlighting the unique impact of N1+SOD on RBC EV characteristics.

### Result 3: Local delivery of RBC EV (N1+SOD) provides protection against retinal degeneration

Given that the combination of N1 and SOD supplementation (N1+SOD) results in a higher RBC EV concentration and distinct proteomic changes indicative of an anti-inflammatory phenotype, we compared four supplementation strategies to identify the most effective *in-vivo* candidate: RBC EV (N1), RBC EV (SOD), RBC EV (N1+SOD), and RBC EV (PBS). The therapeutic efficacy of RBC EV derived from N1 and/or SOD supplementation was investigated in a rodent model of retinal degeneration [47], using intravitreal administration **(Supplementary Figure 5)**. Following 5 days of photo-oxidative damage, retinal function was measured using ERG. Results demonstrated no significant change between the supplementation groups for both a-wave **(Supplementary Figure 5A)** and b-wave **(Supplementary Figure 5B)** responses. However, RBC EV (N1+SOD) showed significantly reduced levels of photoreceptor cell death as measured by TUNEL assay **(Supplementary Figure 5C, P<0.05)**, photoreceptor row counts **(Supplementary Figure 5D, P<0.05)**, and retinal thickness measurements via OCT **(Supplementary Figure 5E-F, P<0.05)**, with representative fundus images **(Supplementary Figure 5G, P<0.05)**.

Therefore, given the proteomics and above *in vivo* findings, the therapeutic efficacy of RBC EV (N1+SOD) was further investigated against a PBS vehicle control (**Figure 3A**), following 5 days of photo-oxidative damage. Results demonstrated that RBC EV (N1+SOD)-injected mice had significantly preserved function for both a-wave (**Figure 3B, P<0.05)** and b-wave (**Figure 3C; P<0.05)** responses compared to PBS-injected controls, along with significantly reduced levels of photoreceptor cell death as measured by TUNEL assay (**Figure 3D, P<0.05)** and photoreceptor row counts (**Figure 3E, P<0.05).** Further, RBC EV (N1+SOD)-injected mice had significantly reduced numbers of IBA-1+ cells in the outer retina, indicating reduced inflammatory cell presence (**Figure 3F; P<0.05)**. Representative confocal images illustrate retinal protection against cell death and inflammation following local administration of RBC EV (N1+SOD) (Figures 3G-H). TUNEL+ cells were also observed in the INL (Figure 3G), with reduced overall INL signal in RBC EV (N1+SOD)-injected retinas compared to the PBS vehicle control, however INL TUNEL+ cells were not quantified in this experiment.

**Figure 3:**
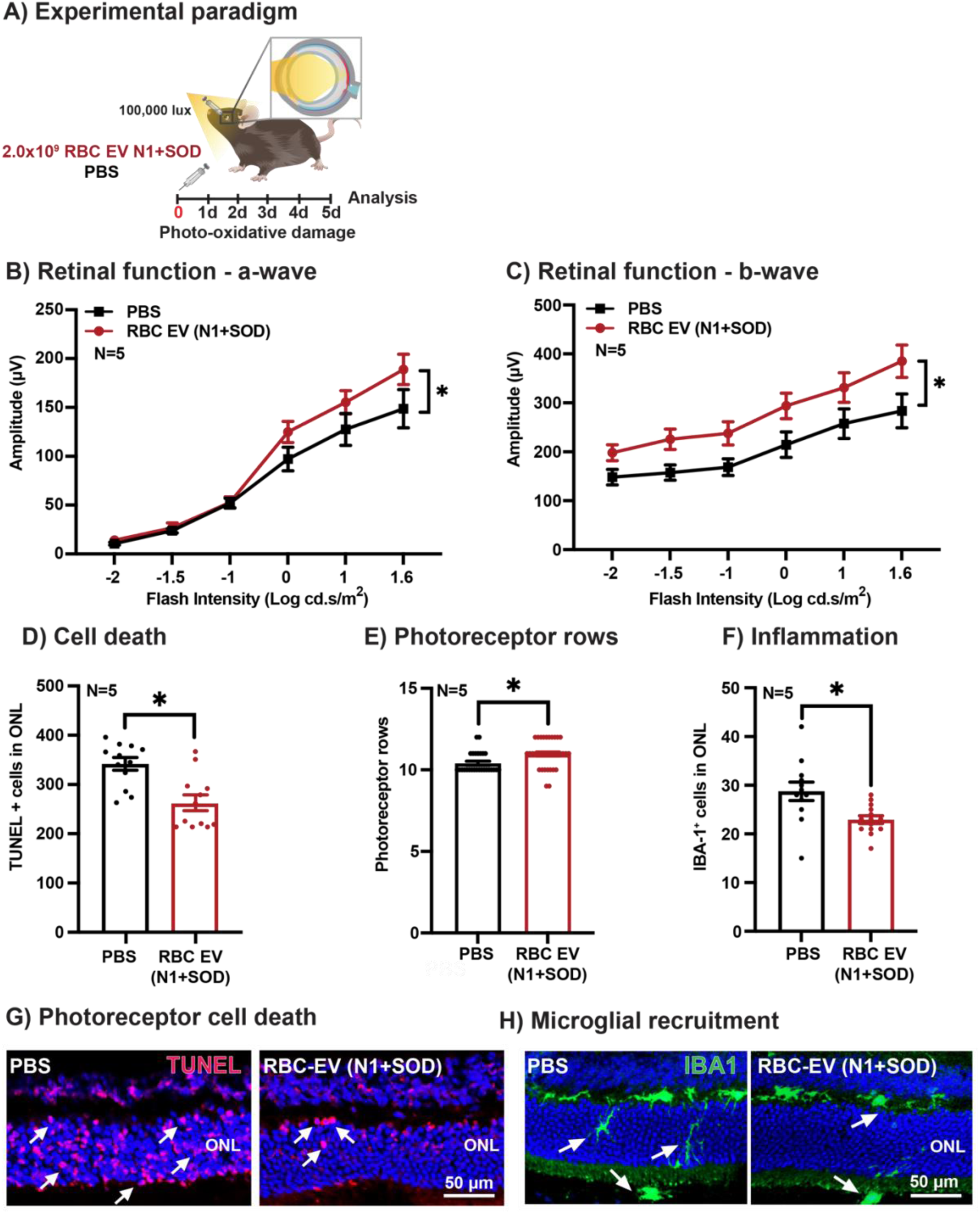
Therapeutic potential of local RBC EV delivery. **(A)** Experimental paradigm for intravitreal injections of RBC EV (N1+SOD). Retinal function as measured by ERG shows **(B)** significantly preserved a-wave function and **(C)** b-wave function compared to PBS controls. **(D)** TUNEL^+^ cells in the ONL were significantly reduced in RBC EV (N1+SOD) compared to controls, while **(E)** photoreceptor preservation was found to be significantly increased in RBC EV (N1+SOD) compared to PBS. **(F)** IBA-1+ microglia were found to be significantly reduced in RBC EV (N1+SOD) mice compared to PBS controls. **(G-H)** Representative confocal images show decreased photoreceptor cell death and the presence of inflammatory cells in the outer retina in RBC EV (N1+SOD) groups. (N=5, P<0.05). Scale = 50μM.

Overall, these results support the protective and anti-inflammatory effects of RBC EV (N1+SOD), highlighting its potential as a therapeutic for retinal degeneration. This therapeutic not only reduces inflammation and cell death but also significantly preserves retinal function.

### Result 4: RBC EV (N1+SOD) is safely taken up across the retina following local delivery

Given the protective effects of N1+SOD supplementation, the safety of RBC EV (N1+SOD) as a local therapeutic was assessed. Mice were injected with RBC EV (N1+SOD) using intravitreal injection and left for 7 days in dim-reared condition (**Figure 4A**). Retinal function was measured using ERG, showing no significant difference in either a-wave (**Figure 4B**) and b-wave responses (**Figure 4C**) compared to PBS injected controls (P>0.05). No differences were observed for measures of cell death (**Figures 4D-E; P>0.05)**, or inflammation (**Figure 4F; P>0.05)**, as measured by TUNEL and IBA1, and shown in representative confocal images (**Figures 4G**). Minimal TUNEL signal was observed in the INL (**Figure 4G),** however INL TUNEL+ cells were not quantified in this experiment. Overall, RBC EV (N1+SOD) did not confer any toxicity or damage to the retina at the parameters measured, suggesting they can be used as a safe therapeutic via local intravitreal injection.

**Figure 4:**
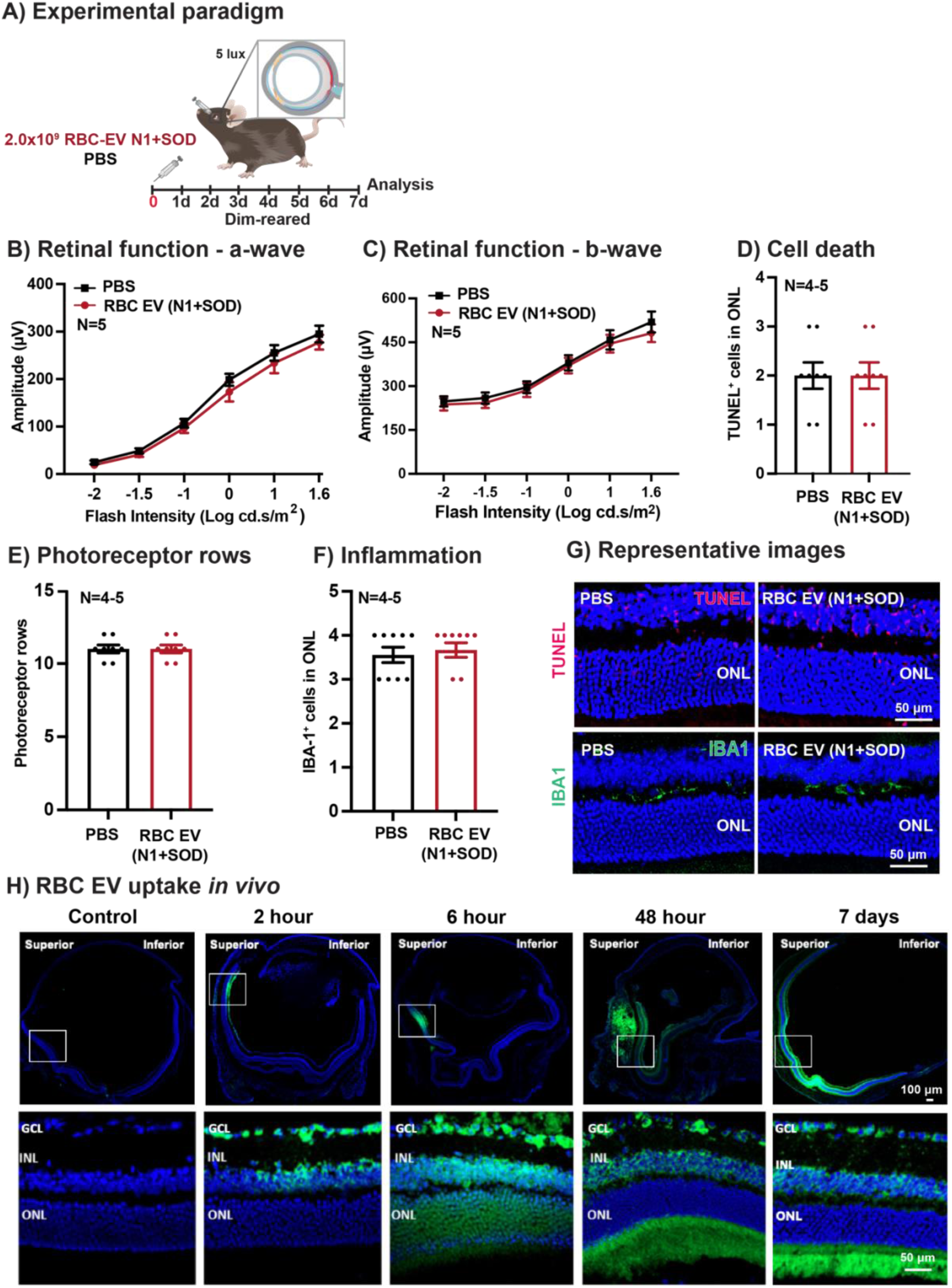
Safety and distribution of local RBC EV (N1+SOD) delivery. **(A)** Experimental paradigm. Retinal function for both **(B)** a-wave and **(C)** b-wave responses was unchanged compared to PBS-injected controls. Further, there were no significant differences in **(D)** TUNEL^+^ cells in the ONL, **(E)** number of photoreceptor rows, or **(F)** presence of IBA-1^+^ microglia in the outer retina. **(G)** Representative confocal images show no differences in TUNEL^+^ cells and IBA-1^+^ cells in outer retina between groups. (N=4-5, P>0.05). Scale = 50μM. **(H)** SYTO-labelled RBC EV (N1+SOD) (green) uptake *in vivo* following intravitreal delivery over 7 days shows progressive radial uptake from the superior retinal delivery site, as well as penetrance through all retinal layers and across the superior and inferior retina by 7 days. (N=1) ONL = outer nuclear layer, INL = inner nuclear layer, GCL = ganglion cell layer. Scale bars = 100μm and 50μm.

In addition, to determine uptake efficiency of RBC EV (N1+SOD) in the retina, RBC EV (N1+SOD) fluorescently labelled with SYTO RNASelect (green) were administered using intravitreal injection. Retinal uptake *in vivo* was observed over 7 days following administration. Uptake could be observed as early as 2 hours post-injection at the site of injection (superior retina) and spread progressively radially across the retina (superior-inferior) as well as penetrating through the retinal layers, with strong labelling observed within the ONL by 6 hours and outer segment labelling from 48 hours. Labelling was still observed at 7 days throughout the retina (**Figures 4H).**

These results support the use of RBC EV (N1+SOD) as a safe and effective therapeutic for retinal degeneration, demonstrating strong uptake and distribution in the retina.

### Result 5-6 : Systemic delivery of RBC EV (N1+SOD) confers retinal protection against photo-oxidative damage and ensures retinal safety

Following local delivery, to assess less-invasive delivery routes, systemic administration was explored for safety and therapeutic efficacy. The therapeutic efficacy of systemic RBC EV (N1+SOD) was investigated using daily intraperitoneal injections at a dose of 2.0×10¹¹ EV/100μL over five days of photo-oxidative damage, compared to PBS control (**Figure 5A).** Retinal function measured using ERG showed significantly preserved retinal function in RBC EV (N1+SOD)-injected mice, for both a-wave (**Figure 5B; P<0.05)** and b-wave (**Figure 5C; P<0.05)** responses. Further, compared to PBS-injected controls, RBC EV (N1+SOD)-injected mice had significantly reduced levels of TUNEL^+^ cells in the ONL (**Figure 5D; P>.0.5)**, significantly higher numbers of photoreceptor rows (**Figure 5E; P<0.05)**, and significantly reduced numbers of IBA-1^+^ microglia/macrophages in the outer retina (**Figure 5F; P<0.05)**. Representative confocal images show decreased levels of cell death and inflammation in the retina of systemically injected RBC EV (N1+SOD) mice (**Figures 5G-H**).

**Figure 5:**
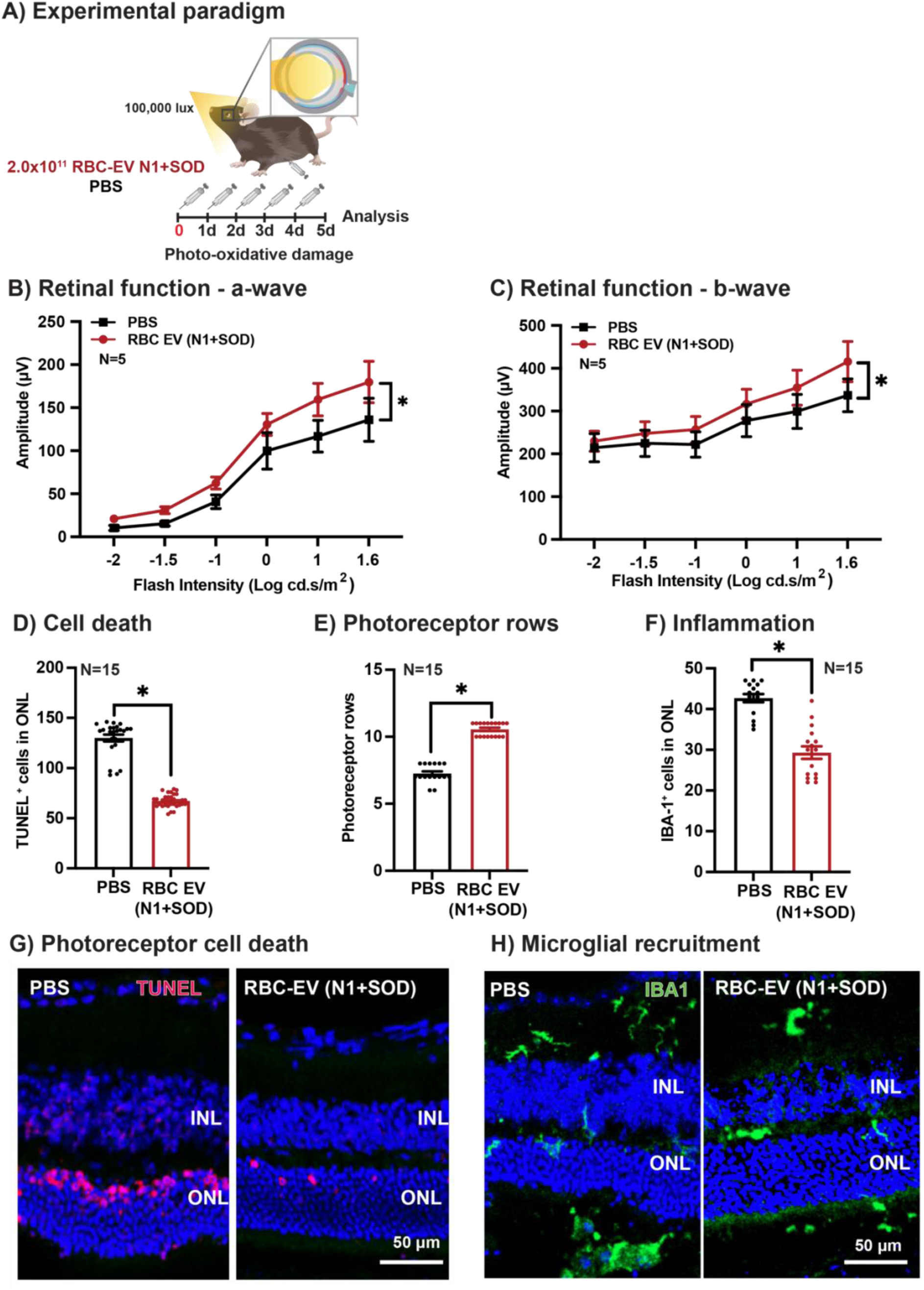
Therapeutic potential of systemic RBC EV (N1+SOD) delivery. **(A)** Experimental paradigm for intraperitoneal injections of RBC EV (N1+SOD). Retinal function as measured by ERG shows **(B)** significantly preserved a-wave function and **(C)** b-wave function compared to PBS controls (N=5, P<0.05). **(D)** TUNEL^+^ cells in the ONL were significantly reduced in RBC EV (N1+SOD) compared to controls, while **(E)** photoreceptor preservation was found to be significantly increased in RBC EV (N1+SOD) compared to PBS. **(F)** IBA-1^+^ microglia were found to be significantly reduced in RBC EV (N1+SOD) mice compared to PBS controls. **(G)** Representative confocal images show reduced TUNEL^+^ cells in the ONL and preserved photoreceptor thickness in RBC EV (N1+SOD)-injected mice and **(H)** reduced IBA-1^+^ cells in the outer retina, compared to PBS. (N=15, P<0.05). ONL = outer nuclear layer. Scale = 50μM.

Similarly, the safety of systemic RBC EV (N1+SOD) administration was investigated by injecting the EVs daily for 7 days via intraperitoneal injection at a dose of 2.0×10^11^ EV/100μL, compared to PBS-injected controls (**Figure 6A).** Following 7 days of dosing under dim-reared condition, retinal function and morphological assessments were conducted. No significant differences were found in retinal function for a-wave (**Figure 6B; P>0.05)** or b-wave (**Figure 6C; P>0.05)** measures between groups. Additionally, there were no differences in levels of cell death (**Figures 6D-E; P>0.05)** or inflammation (**Figure 6F; P>0.05),** as shown in representative confocal images (**Figures 7G-H).**

**Figure 6:**
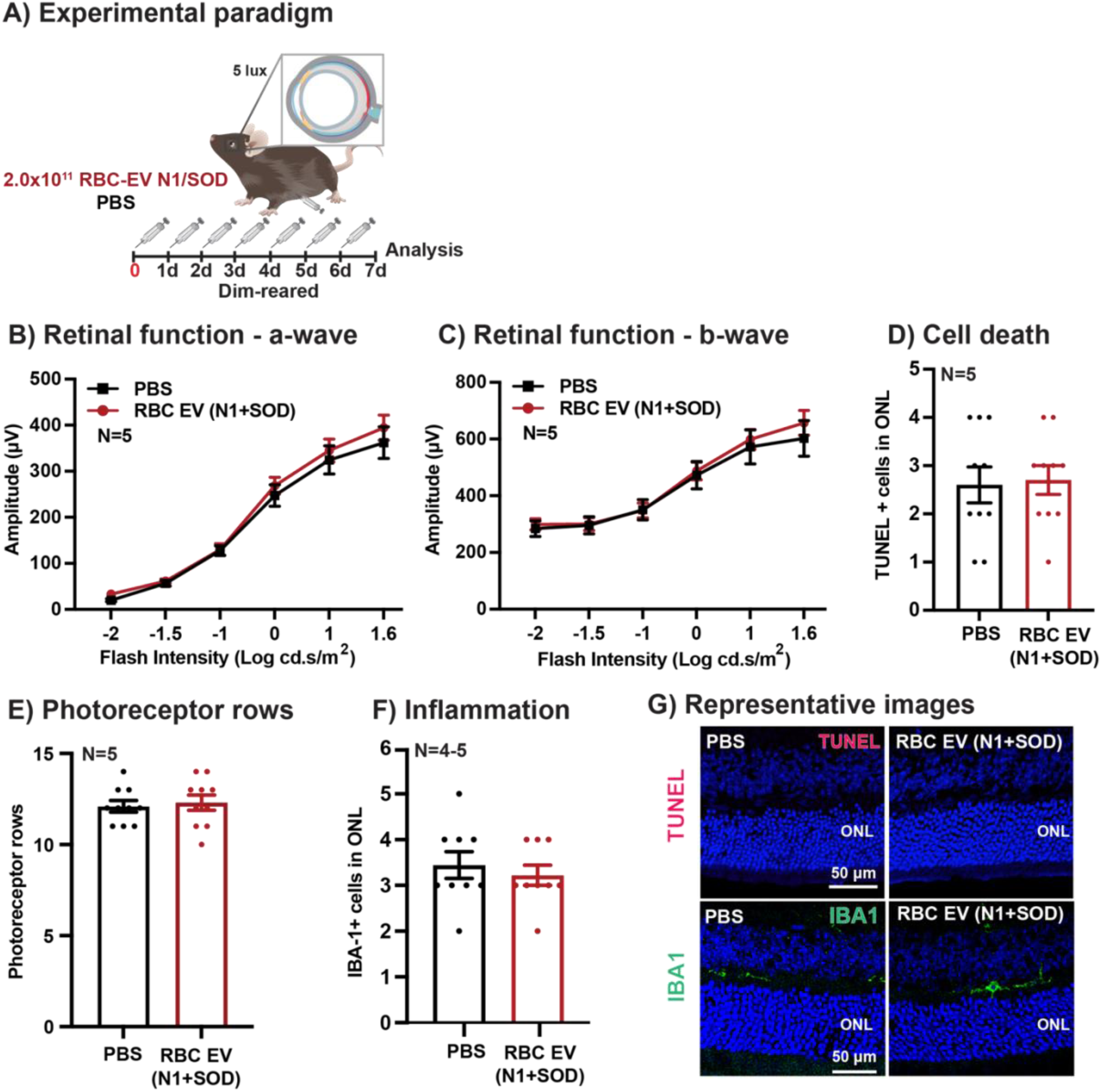
Safety of systemic RBC EV (N1+SOD) delivery. **(A)** Experimental paradigm. Retinal function for both **(B)** a-wave and **(C)** b-wave responses was unchanged compared to PBS-injected controls. Further, there were no significant differences in **(D)** TUNEL^+^ cells in the ONL, **(E)** the number of photoreceptor rows, or **(F)** presence of IBA-1^+^ cells in the outer retina between groups. **(G-H)** Representative confocal images show no differences in TUNEL^+^ cells and IBA-1^+^ cells between groups. (N=4-5, P>0.05). ONL = outer nuclear layer. Scale = 50μM.

**Figure 7:**
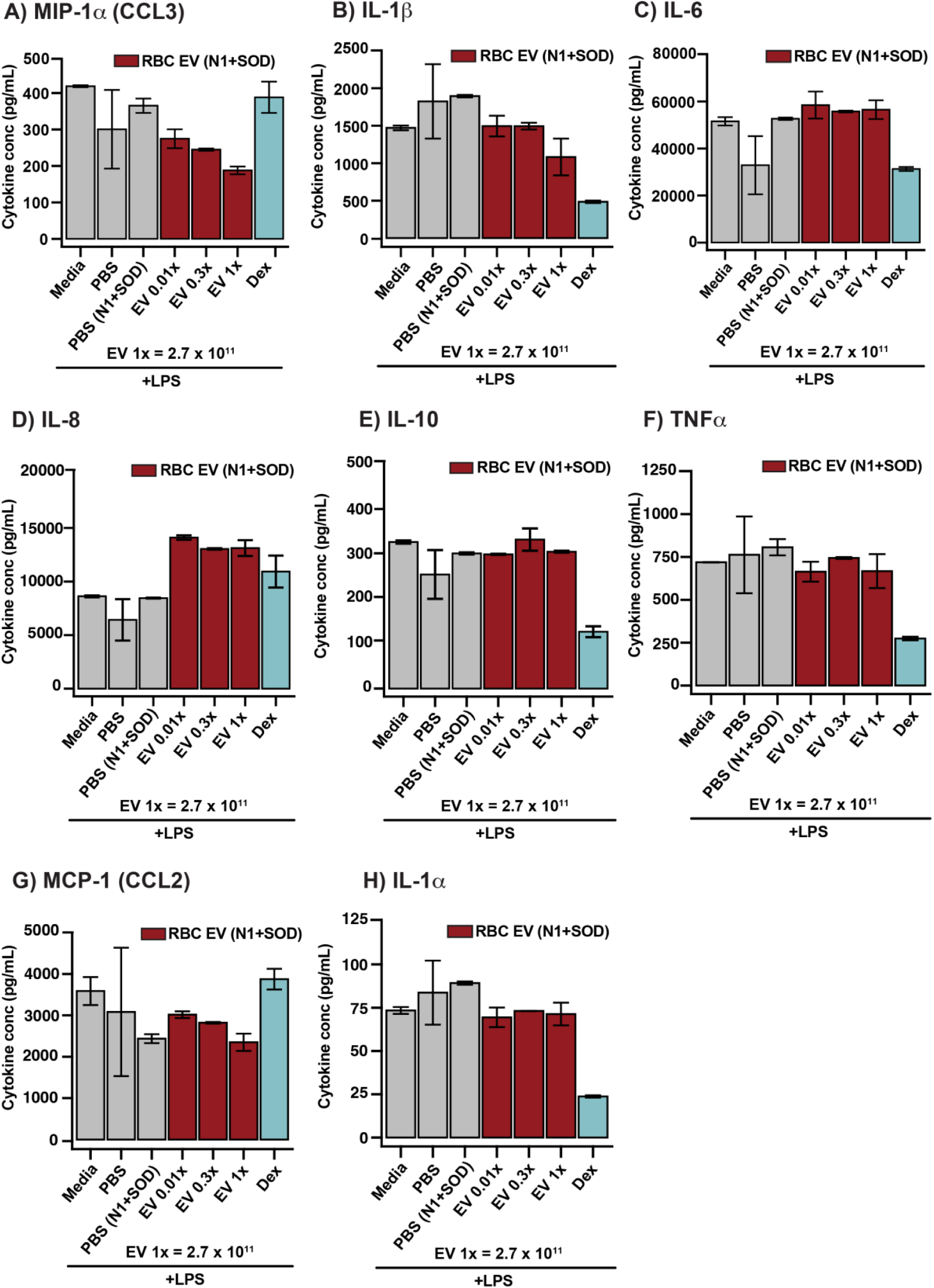
RBC EV (N1+SOD) decrease key pro-inflammatory cytokine production in PBMC. **(A-H)** Cytokine output following incubation of LPS-stimulated PBMCs with RBC EV (N1+SOD) at 1x, 0.3x and 0.01x at 1x=2.7 x 10^11^, for 48 hours, N=1, run in duplicates.

In conclusion, systemic administration of RBC EV (N1+SOD) demonstrates a promising therapeutic potential for protecting the retina against degeneration. The treatment did not exhibit any signs of toxicity to the retina after 7 days of daily dosing, suggesting it is a safe and effective approach. These findings further support the investigation and development of RBC EV (N1+SOD) as a viable therapy for retinal protection.

### Result 7: RBC EV (N1+SOD) have anti-inflammatory properties and can reduce inflammatory cytokine release from PBMC

Finally, in order to assess and verify the observed immuno-modulatory properties of RBC EV (N1+SOD) identified from multi-omic analyses and *in vivo* studies, RBC EV (N1+SOD) were sent for independent validation for measures of metabolic activity, safety, and anti-inflammatory properties. RBC EV (N1+SOD) at 3 concentrations (1x, 0.3x and 0.01x, 1x=2.7 x 10^11^) were incubated on peripheral bone mononuclear phagocyte cells (50,000 PBMC/well) with inflammatory stimulation (LPS at 100 ng/mL). Further, dose responses were also evaluated. Cytokine outputs and viability measures were compared to control groups - PBS, PBS (N1+SOD), media, and dexamethasone 10nM.

Results suggested a dose-dependent increase in PBMC metabolic activity in response to increased RBC EV (N1+SOD) concentration in both control **(Supplementary Figure 6A)** and LPS-stimulated cells **(Supplementary Figure 6B)**.

RBC EV (N1+SOD) were also assessed for their safety profile on PBMCs, with cytokine output MIP-1α (CCL3), IL-1β, IL-6, IL-8 IL-10, TNFα, MCP-1 (CCL2) and IL-1α measured following 48 hours incubation **(Supplementary Figure 7).** These cytokines play known pathogenic roles in both retinal and neurodegenerative diseases [55–57]. While RBC EV (N1+SOD) increased basal inflammatory cytokine release over media control in the unstimulated condition, this was comparable to PBS and/or below the LLOQ (lower limit at which assay can provide quantitative results), with the exception of MCP-1 (CCL2) at 0.3x dose **(Supplementary Figure 7G)**. Overall, these results support broad safety of RBC EV (N1+SOD).

Following this, RBC EV (N1+SOD) were assessed for their anti-inflammatory properties using the same cytokine outputs and following 48 hours LPS stimulation and RBC EV (N1+SOD) incubation (**Figure 7A-H**). Results showed an anti-inflammatory response for cytokines MIP-1α (CCL3), IL-1β and MCP-1 (CCL2), **(Figure 7A, B and G)**, with a dose-dependent decrease for cytokines MIP-1α (CCL3) and MCP-1 (CCL2). Importantly, RBC EV (N1+SOD) was able to reduce cytokine output below levels of anti-inflammatory steroid dexamethasone (10nM) for MIP-1α (CCL3) and MCP-1 (CCL2) (**Figure 7 A and G)**.

Taken together, these results collectively support the therapeutic use of RBC EV (N1+SOD) in reducing neuroinflammation and cell death, and in preserving retinal function through both local and systemic administration. The strong mechanism of action for RBC EV (N1+SOD) observed through omics, along with their ability to reduce release of key pro-inflammatory cytokines involved in retinal and neurodegenerative diseases, further highlights their potential as neuroinflammatory modulators.

## Discussion

This study highlights the potential therapeutic application of RBC EV to mediate neuroinflammatory responses associated with retinal degenerations and suggests their broad applicability for treating retinal and neurodegenerative diseases where neuroinflammation is an underlying disease feature. We demonstrated that N1+SOD supplementation significantly influenced an anti-inflammatory RBC profile, resulting in a high-yield, reproducible, and neuroprotective RBC EV population. *In vivo*, local, and systemic administration of RBC EV (N1+SOD) provided significant protection against retinal degeneration, as demonstrated by significantly preserved retinal function, increased photoreceptor survivability, and decreased inflammation, while being safe with no signs of retinal toxicity. Additionally*, in vitro* cytokine assays strongly supported the anti-inflammatory mechanism-of-action observed *in vivo*, with RBC EV (N1+SOD) reducing the production of known pro-inflammatory cytokines involved in AMD and broader neurodegenerative diseases. Altogether, these findings demonstrate the use of RBC EV to modulate neuroinflammation and supports their broad applicability and exciting translational potential to be developed as less-invasive therapeutics for use in the CNS.

### RBC EV (N1+SOD) are ideal therapeutics for treating neuroinflammation

Neuroinflammation is a hallmark feature of retinal and neurodegenerative diseases, characterised by recruitment and activation of resident and peripheral immune cells, and pro-inflammatory cytokine release leading to neuronal cell death [2]. For the majority of retinal and neurodegenerative diseases, there are few, to no, treatments available. Given the significant contribution of inflammation to disease onset and progression, targeting inflammatory pathways to reduce the severity of disease is therefore a viable therapeutic strategy.

RBC EV have garnered significant interest as a lowly-immunogenic, and biologically compatible therapeutic with results from this work and others supporting their use in treating chronic inflammation [35–37, 40, 58]. Unlike stem cells and other primary cells in the body, as RBC do not have nuclear or mitochondrial DNA, RBC-EV have reduced molecular cargo limiting horizontal transfer and adverse effects in recipient cells and the development of oncogenic phenotypes [36, 43, 59, 60]. As the most abundant and non-inflammatory cell in the body, RBC can be readily collected and processed on a large scale for non-immunogenic EV, overcoming challenges of high costs and scalability associated with culturing and maintaining primary cells for EV isolation [41, 61]. By utilising the body’s own cells and natural communication pathways, these natural drug delivery vehicles can be readily collected from both patients and donors and redelivered as a safe autologous/allogenic therapeutic, improving patient safety and reducing adverse effects as a personalised treatment. In fact, RBC EV have previously demonstrated anti-inflammatory benefits in treating atherosclerosis, [40] and have shown therapeutic effects in various cancers, including metastatic breast cancer [36] and in models of cancer cachexia [37].

To improve upon the inherent protective qualities of RBC EV, in this work we showed that RBC EV (N1+SOD) derived using our novel incubation pipeline exhibited a unique anti-inflammatory and therapeutic profile compared to baseline RBC EV (PBS), or other supplementation conditions tested. The proteomic profile of RBC EV (N1+SOD) showed significant downregulation in proteins such as CRLF3, TOLLIP, EIF5, SNX15, WNK1 and GD1, which are involved in the transcription of pro-inflammatory genes, inflammasome-associated pathways, and interleukin-1 signalling [62–66]. This anti-inflammatory proteomic profile was further supported by anti-inflammatory microRNA cargo, including miR-451a and miR-let-7i-5p, which are known inhibitors of the TLR4/Nfκ-B inflammatory cascade [67]. This aligns with the significant therapeutic advantage of RBC EV (N1+SOD) *in vivo* over RBC EV (PBS) and other supplementation conditions in reducing neuroinflammation and neuronal cell death in the retina. We attribute this unique anti-inflammatory profile and improved protection against neuroinflammation to our incubation pipeline, which includes N1 medium components and superoxide dismutase (SOD). N1 medium components-transferrin, has previously been shown to reduce reactive oxidant species (ROS) formation [68], and progesterone, which conserves photoreceptors and reduces cell death in models of retinal degeneration along with SOD, which breaks down superoxide radicals; collectively enhancing neuroprotection [69]. We hypothesise that this combined approach mitigates excess ROS production, thereby protecting RBC during the incubation process to obtain high quality neuroprotective and anti-inflammatory EV, and to the retina during therapeutic delivery supporting RBC EV (N1+SOD) as an ideal therapeutic for treating neuroinflammation in retinal degenerations.

### RBC EV (N1+SOD) reduced neuroinflammation in the retina

We showed in this work that RBC EV (N1+SOD) were able to modulate broad neuroinflammation by regulating the release of multiple pro-inflammatory cytokines, addressing the need for multi-pathway modulation in these diseases. The marked reduction in pro-inflammatory cytokine levels - MIP-1α (CCL3), IL-1β, and MCP-1 (CCL2), is particularly significant as they play key roles in the onset and progression of various retinal and neurodegenerative conditions [12, 47, 54, 70–75]. MIP-1α (CCL3), expressed by astrocytes, glia, and neurons in the CNS, recruits circulating inflammatory cells to sites of damage, exacerbating inflammation and leading to neurodegeneration, including conditions like ischemic stroke, Alzheimer’s disease, and various retinal degenerations such as neovascularization, ischemic retinopathy, and AMD [70, 71]. Similarly, IL-1β, a pleiotropic cytokine upregulated in Alzheimer’s disease, Parkinson’s disease, and AMD, activates microglia and astrocytes, stimulating the production of downstream pro-inflammatory cytokines [76]. Additionally, CCL2 controls the migration of monocytes/macrophages and dendritic cells [71, 74], and is involved in neurodegenerative disorders, including Alzheimer’s disease, MS, and is significantly elevated in patients with both wet and dry AMD [47]. Our previous work has shown that reducing the expression of CCL3, CCL2, and IL-1β in the retina suppresses macrophage accumulation, decreases retinal inflammation and cell death [77, 78], supporting the protective mechanism of action of RBC EV (N1+SOD) observed *in vivo.* Furthermore, cytokines CCL2 and IL-1β are significantly upregulated in the early stages of retinal degeneration, preceding the onset of photoreceptor cell death [3], highlighting the potential of RBC EV (N1+SOD) as an early intervention for mitigating neuroinflammation and preserving tissue health before the onset of irreversible neuronal cell death in both retinal and neurodegenerative diseases.

### Mechanism of systemic RBC EV (N1+SOD) administration for modulating neuroinflammation

Current challenges in developing therapeutics for combating neuroinflammation in the CNS include overcoming blood-tissue barriers including the BRB and BBB, which exclude most therapeutics following systemic administration. This challenge often necessitates more invasive delivery methods such as intravitreal delivery for retinal degenerations, or intracranial for neurodegenerations, both requiring specialised clinical care, and limiting the number of patients that can be treated due to a bottleneck in available specialists and higher costs [79]. RBC EV are a promising alternative, evidenced from the significant anti-inflammatory therapeutic effects seen when delivered systemically in this work.

We propose that systemic administration of RBC EV (N1+SOD) could provide therapeutic effects in the retina and wider CNS through several mechanisms. Firstly, EV have been shown to target and accumulate at sites of vascular damage [80], degeneration [81], or compromised blood-tissue barriers [82]. In the photo-oxidative damage model of retinal degeneration, which features compromised blood-tissue barriers and retinal cell death, this targeting ability suggests that anti-inflammatory RBC EV (N1+SOD) could provide localised protection to the retina following systemic administration. Secondly, RBC EV naturally target granulocytes, monocytes, and endothelial cells, which are crucial in maintaining the innate immune system within the CNS, which plays a key role in both retinal- and neurodegeneration [83]. RBC EV (N1+SOD) may target the innate immune system and circulating immune cells, preventing their infiltration into the retina and thereby mitigating degeneration. Additionally, RBC contain key complement-regulatory proteins, including CD55 and CD59, with previous studies showing AAV-mediated delivery of soluble CD59 (a C9 inhibitor) able to reduce membrane attack complex and NLRP3 inflammasome activation [84], both critical components of the innate immune system implicated in AMD [83]. Finally, as most EV types, including RBC EV, accumulate in the liver when delivered systemically, we hypothesise that local accumulation could provide anti-inflammatory effects due to the presence of phosphatidylserine (PS) on the EV plasma membrane [40]. PS, which is essential for neuron survival [85], has been previously shown to alter levels of pro-inflammatory cytokines in the brain when administered systemically, thereby boosting the central immune system against inflammatory insults [86]. In addition, as complement proteins are produced by the liver, we also hypothesise that systemic inhibition of liver-produced complement components may modulate circulating inflammation levels and induce a subsequent reduction in local retinal inflammation during degeneration [87, 88]. This hypothesis is supported from works showing the effects of liver produced complement proteins on inducing local complement activation in the eye [89]. The future development of this less-invasive approach would be a game-changer in the ophthalmic and neurodegenerative fields, reducing associated costs and increasing treatment accessibility to the wider population.

## Conclusion

Neuroinflammation in the CNS significantly contributes to neurodegeneration, making it a critical target for therapeutic interventions aimed at treating retinal- and neurodegenerative diseases. Challenges in developing therapeutics for neuroinflammation include overcoming blood-tissue barriers and modulating multiple neuroinflammatory pathways within immune-privileged environments. This study highlights the potential of RBC EVs (N1+SOD) in overcoming these challenges and effectively mediating neuroinflammation. We demonstrate that *in vivo* administration of RBC EVs (N1+SOD), both locally and systemically, preserved retinal function, increased photoreceptor survival, and reduced inflammation in a murine model of retinal degeneration. *In vitro* assays further support these neuro-modulatory effects by reducing pro-inflammatory cytokine production. These findings support the use of RBC EV (N1+SOD) for the treatment of retinal degenerations and broader neurodegenerative diseases, offering a scalable, effective, and less-invasive systemic treatment approach.

## Author contributions

Rakshanya Sekar, Adrian V. Cioanca, Riccardo Natoli, and Yvette Wooff: conceptualization. Rakshanya Sekar, Adrian V. Cioanca, Yilei Yang, Karthik Kamath, Luke Carroll, and Yvette Wooff: methodology and data analysis. Rakshanya Sekar, Adrian V. Cioanca and Yvette Wooff: investigation. Rakshanya Sekar and Yvette Wooff: writing—draft, review, and editing. Riccardo Natoli and Yvette Wooff: funding acquisition. This research used NCRIS-enabled Australian Proteome Analysis Facility (APAF) infrastructure.

## Acknowledgements

An acknowledgement is made to the donors of the Macular Disease Research, a program of the BrightFocus Foundation (M2021012F), and to the Macular Disease Foundation Australia (MDFA), for support of this research awarded to Yvette Wooff, and to the ANU Impact Fund (IF601) and The National Health and Medical Research Council of Australia (NHMRC: 202239) awarded to Riccardo Natoli.

## List of Abbreviations

AEC: Animal Ethics Committee
AMD: Age related macular degeneration
cDNA: complementary DNA
CO2: Carbon dioxide
DR: Dim-reared
ERG: Electroretinography
EV: extracellular vesicle
HTS: high-throughput sequencing
INL: inner nuclear layer
LED: light-emitting diode
ONL: outer nuclear layer
PBS: phosphate buffered solution
PCA: principal component analysis
PD: Photo-oxidative damage
qPCR: quantitative polymerase chain reaction
RNA: ribonucleic acid
RT: room temperature
RBC EV: Red blood cell derived extracellular vesicles
SOD: Superoxide Dismutase
UMAP: Uniform Manifold Approximation and Projection
WT: Wild Type

