## Supplementary Figures for "Therapeutic potential of red blood cell-derived extracellular vesicles in reducing neuroinflammation and protecting against retinal degeneration"

#reflects equal first co-authorship

*reflects equal senior co-authorship

Corresponding author:

### Supplementary Figure 1

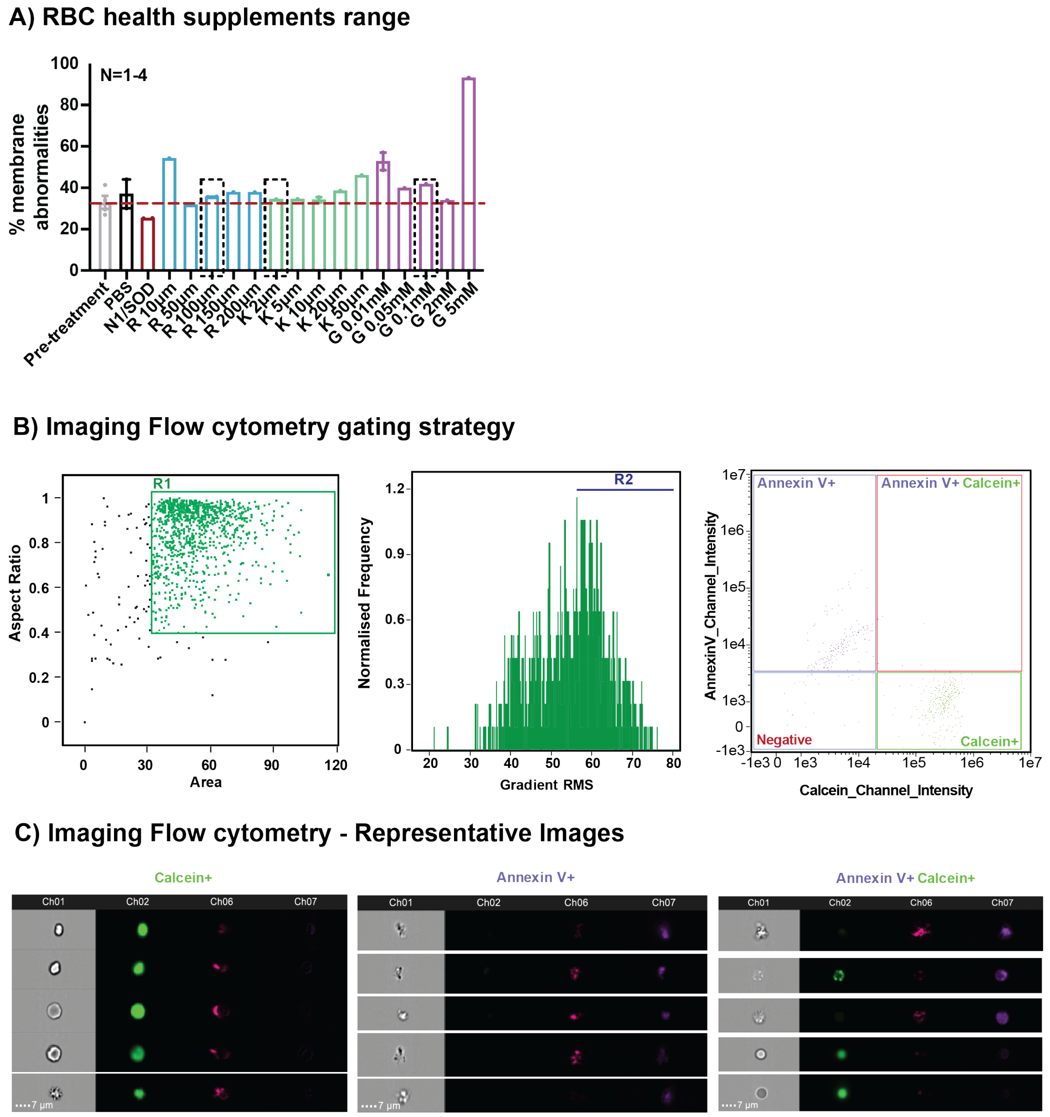


**Supplementary Figure 1: RBC characterisation post supplementation. (A)** Compared to pre-treatment RBC, RBC (N1+SOD) had the lowest % abnormality of all antioxidants and doses tested. RBC with Resveratrol 100uM, Kaempferol (2uM) and Glutathione (0.1mM) had the lowest % abnormality out of the respective dose ranges tested (dashed lines), N=1-4 **(B)** Gating strategy used for Amnis II Imaging flow cytometry, Cells in R2 were characterised as Calcein$^+$, Annexin$^+$ or Double positive- Calcein$^+$Annexin$^{+.}$ **(C)** Representative images show cells in respective gates- Calcein$^+$, Annexin$^+$ and Double positive- Calcein$^+$Annexin$^+$

### Supplementary Figure 2

A) Top 50 Proteins- PBS vs Pretreatment

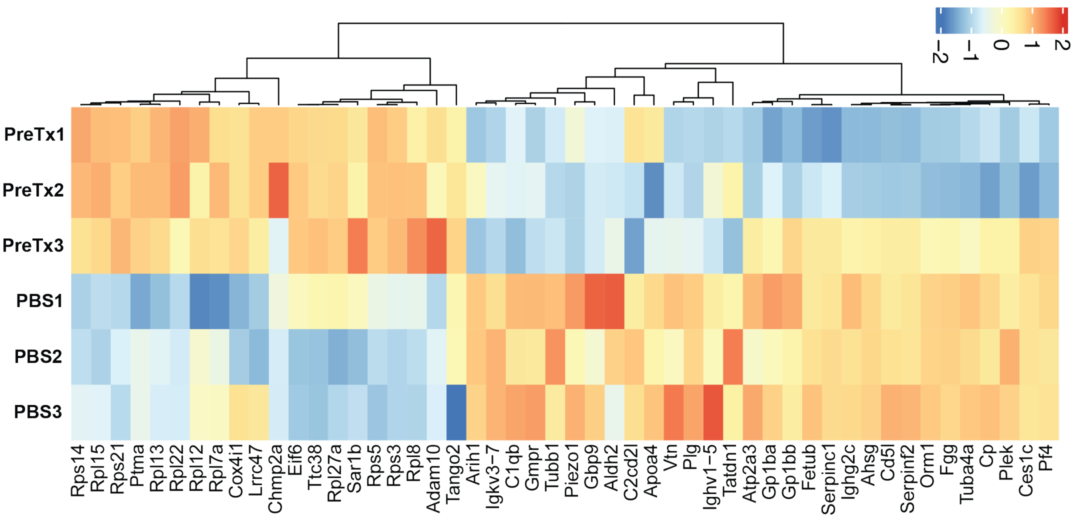


B) Top 20 Pathways

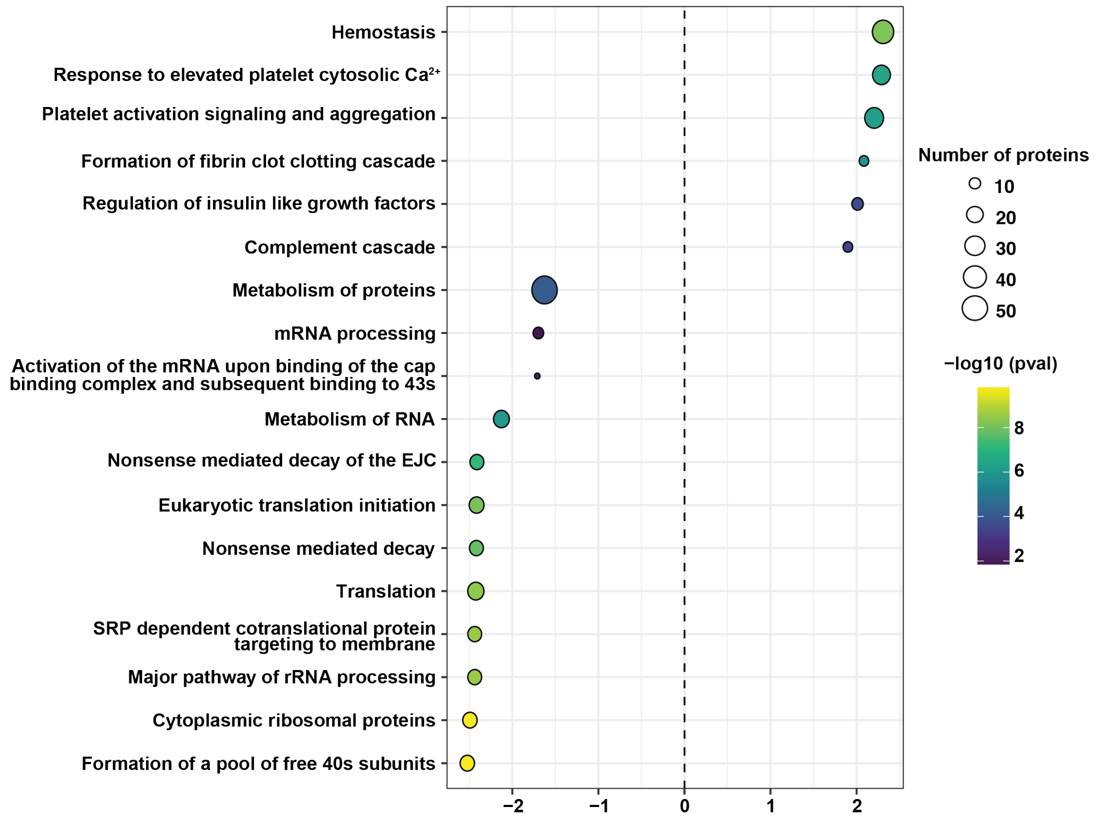


**Supplementary Figure 2: Characterisation of RBC proteomic content post supplementation.** (A) Top 50 differentially expressed proteins between PBS and Pre-treatment. (B) Top 20 pathways, (N=3,$p<0.05$ )

**<u>Supplementary Figure 3</u>**

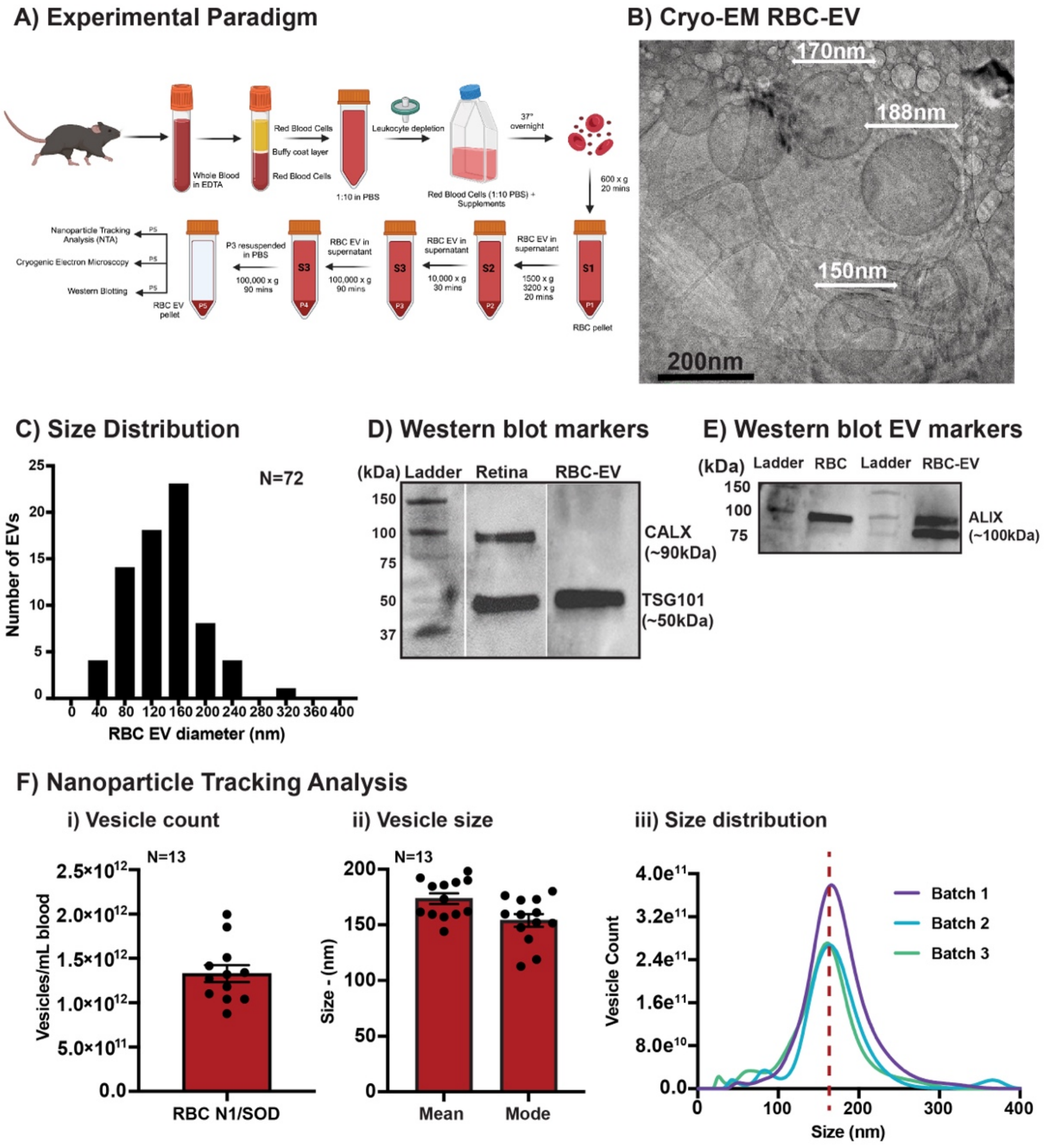


**Supplementary Figure 3: RBC EV characterisation. A)** Experimental paradigm for RBC EV isolation and incubation conditions. **(B)** Cryo-EM imaging of RBC EV (N1+SOD) post isolation. **(C)** Size distribution of RBC EV from cryo-EM images (N=72, Scale = 200nm). **(D-E)** Western blots for cellular marker Calnexin (CNX), EV markers Tumour susceptibility gene 101 (TSG101) and ALIX. (D) CNX (90kDa) band was present in retinal lysates, but not RBC EV lysates, TSG101 was present in both retinal and RBC EV lysates **(E)** ALIX (100kDa) was enriched in RBC EV lysates over RBC lysates.

### Supplementary Figure 4

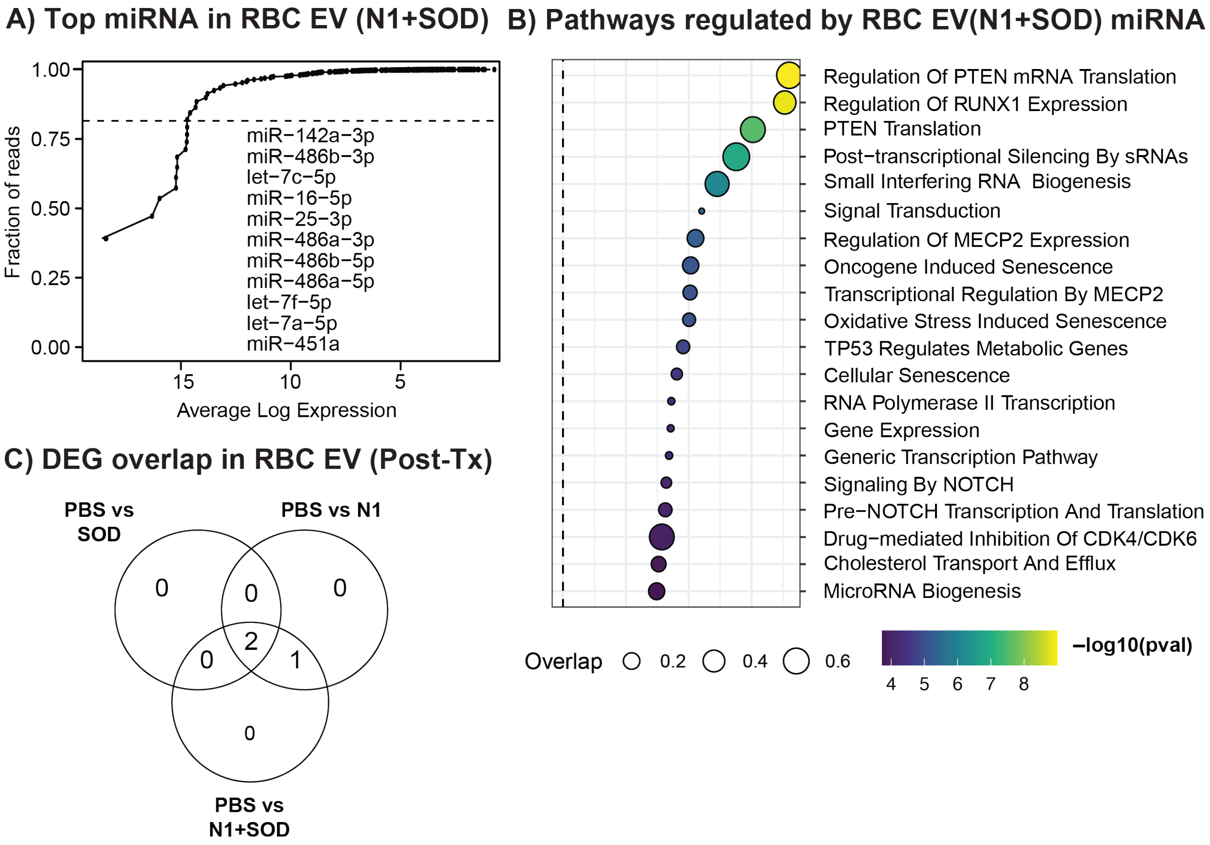


**Supplementary Figure 4: Media supplementations did not change microRNA signature of RBC EV. (A)** Top 11 microRNA abundance. **(B)** Pathway analysis of enriched microRNA. **(D)** Venn diagram showing no unique microRNA between supplement groups. (N=3, P<0.05)

**Supplementary Figure 5**

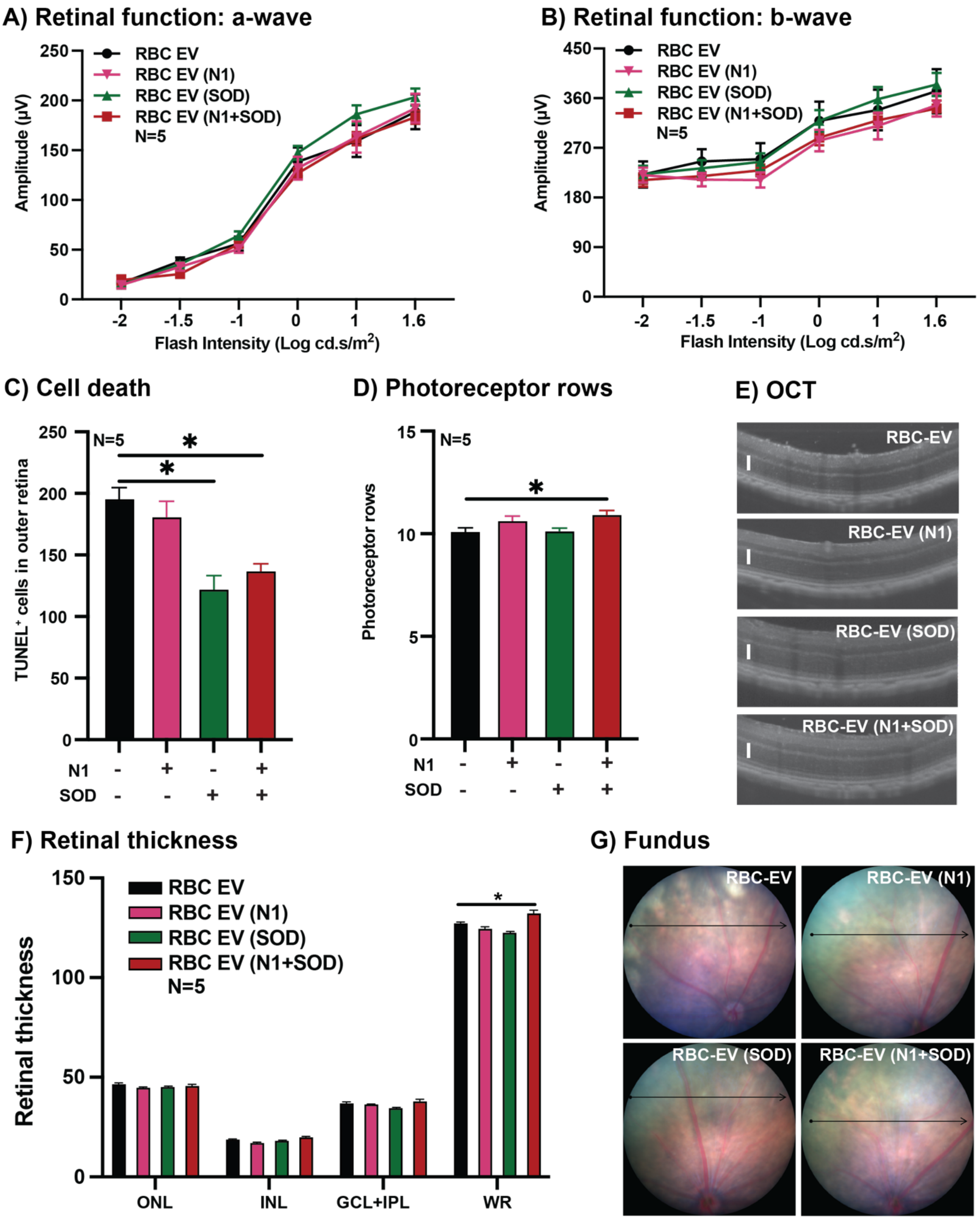

**Supplementary Figure 5: Local delivery of RBC EV via Intravitreal injection. (A)** ERG shows no change in a-wave and **(B)** b-wave function ($P>0.05$, $N=5$). **(C)** TUNEL$^{+}$ cells in the ONL were significantly reduced in RBC EV (SOD) and RBC EV (N1+SOD) groups compared to RBC EV and RBC EV (N1) group, while **(D)** photoreceptor preservation was found to be significantly increased in RBC EV (N1+SOD) group. **(E)** OCT images show improved retinal health in RBC EV (N1+SOD) groups quantified **(F)** Whole retinal thickness was significantly increased between RBC EV and RBC EV (N1+SOD) groups ($P<0.05$, $N=5$). Representative **(G)** fundus images.

**Supplementary Figure 6**

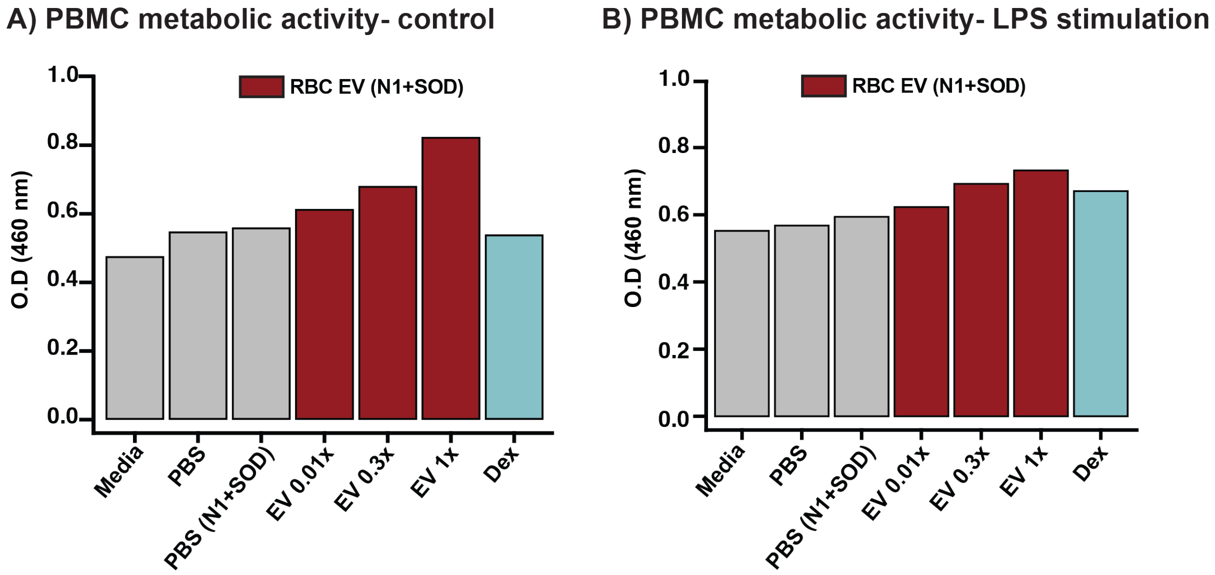


**Supplementary Figure 6: RBC EV (N1+SOD) promoted dose-dependent increase in PBMC metabolic activity.** A dose-dependent response of increased PBMC metabolic activity was shown following RBC EV (N1+SOD) incubation in both **(A)** control and **(B)** LPS-stimulated conditions. N=1, run in biological duplicates.

**Supplementary Figure 7**

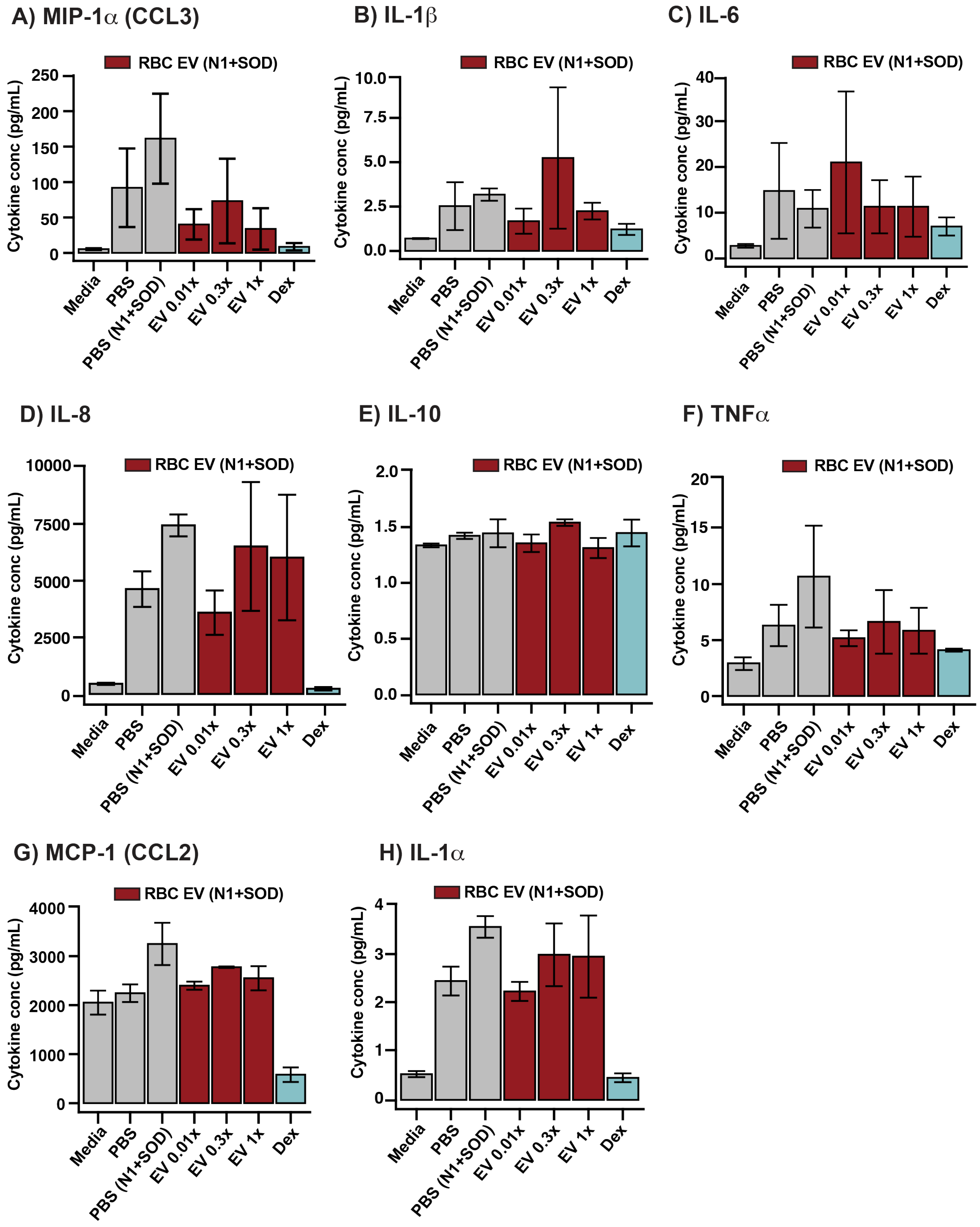


**Supplementary Figure 7: RBC EV (N1+SOD) induce basal inflammatory responses via select pro-inflammatory cytokine. (A-H)** Cytokine output following incubation of PBMCs with RBC EV (N1+SOD) for 48 hours. N=1, run in biological duplicates.

**Supplementary Figure 8**

**A) Full length western blot - CANX and TSG101**

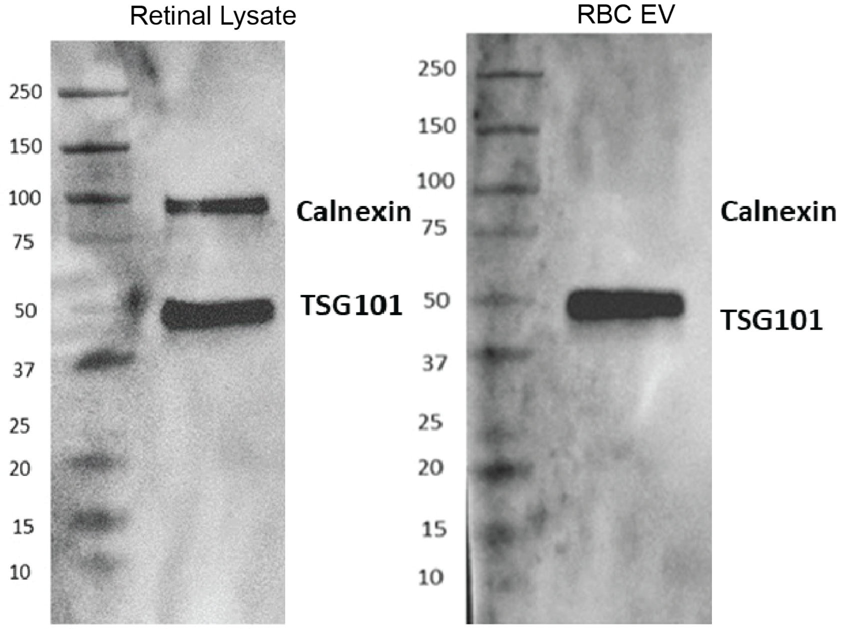


**B) Full length western blot- ALIX**

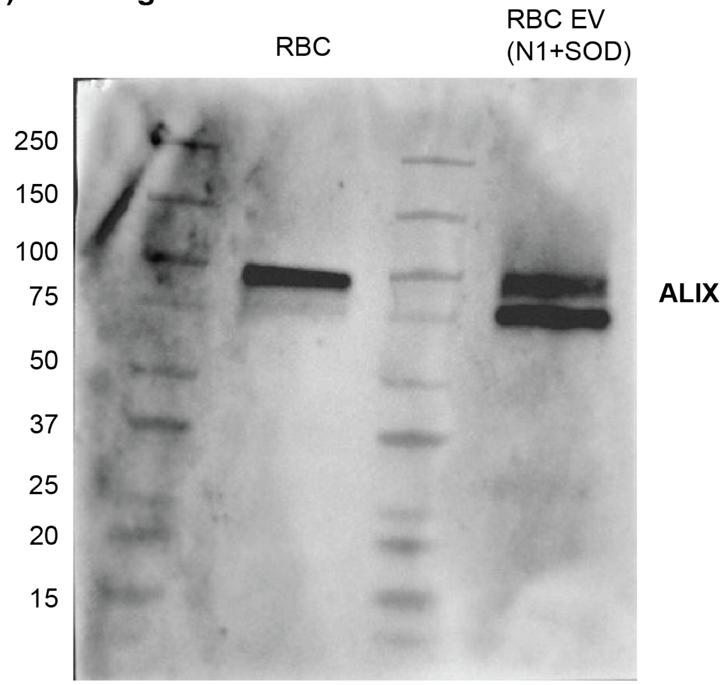


**Supplementary Figure 8: Full western blots showcased in supplementary figure 3. (A)** Retinal protein lysate and RBC EV for Calnexin and TSG101 **(B)** RBC and RBC EV lysate for EV protein ALIX.
